# All-optical electrophysiology with improved genetically encoded voltage indicators reveals interneuron network dynamics in vivo

**DOI:** 10.1101/2021.11.22.469481

**Authors:** He Tian, Hunter C. Davis, J. David Wong-Campos, Linlin Z. Fan, Benjamin Gmeiner, Shahinoor Begum, Christopher A. Werley, Gabriel B. Borja, Hansini Upadhyay, Himali Shah, Jane Jacques, Pojeong Park, Yitong Qi, Vicente Parot, Karl Deisseroth, Adam E. Cohen

**Affiliations:** Department of Chemistry and Chemical Biology, Harvard University, Cambridge, MA, USA; Department of Bioengineering, Department of Psychiatry and Behavioral Sciences, Stanford University, Stanford, CA, USA; Q-State Biosciences, Cambridge, MA, USA; Howard Hughes Medical Institute; Department of Physics, Harvard University, Cambridge, MA, USA

## Abstract

All-optical electrophysiology can be a powerful tool for studying neural dynamics *in vivo*, as it offers the ability to image and perturb membrane voltage in multiple cells simultaneously. The “Optopatch” constructs combine a red-shifted archaerhodopsin (Arch)-derived genetically encoded voltage indicator (GEVI) with a blue-shifted channelrhodopsin actuator (ChR). We used a video-based pooled screen to evolve Arch-derived GEVIs with improved signal-to-noise ratio (QuasAr6a) and kinetics (QuasAr6b). By combining optogenetic stimulation of individual cells with high-precision voltage imaging in neighboring cells, we mapped inhibitory and gap junction-mediated connections, *in vivo*. Optogenetic activation of a single NDNF-expressing neuron in visual cortex Layer 1 significantly suppressed the spike rate in some neighboring NDNF interneurons. Hippocampal PV cells showed near-synchronous spikes across multiple cells at a frequency significantly above what one would expect from independent spiking, suggesting that collective inhibitory spikes may play an important signaling role *in vivo*. By stimulating individual cells and recording from neighbors, we quantified gap junction coupling strengths. Together, these results demonstrate powerful new tools for all-optical microcircuit dissection in live mice.

## Introduction

A central goal of systems neuroscience is to understand the relation between neural circuitry and network dynamics. Experiments to map such relations involve both readout and precise manipulation of circuit dynamics *in vivo*. All-optical electrophysiology represents an attractive approach for this purpose as it combines the power of voltage imaging and optogenetic stimulation (Adam et al., 2019; Emiliani et al., 2015; Fan et al., 2020; Hochbaum et al., 2014; Piatkevich et al., 2019). Voltage imaging with genetically encoded voltage indicators (GEVIs) enables repeated optical readout of subthreshold and spiking membrane voltage dynamics *in vivo* from genetically defined neurons at millisecond temporal resolution (Abdelfattah et al., 2019; Abdelfattah et al., 2021; Adam et al., 2019; Böhm et al., 2021; Chien et al., 2021; Evans et al., 2021; Fan et al., 2020; Gong et al., 2015; Kannan et al., 2021; Marshall et al., 2016; Piatkevich et al., 2019; Villette et al., 2019). Optogenetic perturbations, meanwhile, probe causal relationships in circuit dynamics and behaviors through spatially and temporally precise perturbation of the circuit (Carrillo-Reid et al., 2019; Jennings et al., 2019; Marshel et al., 2019; Packer et al., 2015; Robinson et al., 2020; Wolff et al., 2018).

All-optical electrophysiology requires voltage imaging and optogenetic manipulation to be crosstalk-free. Since all known channelrhodopsins have action spectra which maintain substantial photocurrents in the blue part of the spectrum, crosstalk is minimized by combining a blue-shifted channelrhodopsin with a voltage indicator excited at 590 nm or longer wavelengths. Thus far, only the combination of far-red archaerhodopsin (Arch)-derived GEVIs with blue-shifted channelrhodopsins has demonstrated sufficient spectral orthogonality to meet this standard. However, the *in vivo* performance of Arch-derived GEVIs has been a major challenge for widespread applications of all-optical electrophysiology. Development of improved GEVIs has been limited by the difficulties associated with multi-dimensional functional screening in mammalian cells. Here we developed a novel approach for directed evolution of genetically encoded biosensors using a video-based pooled high-throughput screen. We used this platform to evolve Arch-derived GEVIs with improved signal-to-noise ratio (QuasAr6a) and kinetics (QuasAr6b).

Based on these improved GEVIs, we developed new applications of all-optical electrophysiology to study interneurons in live mice. Inhibitory interneurons play a vital role in maintaining the excitation/inhibition balance within the brain. While interneurons constitute a numerical minority among the entire neuronal population, they can be classified into an expanding list of subtypes based on morphology, electrical properties, connectivity, biomarker expression, and functional role (Lim et al., 2018; Pelkey et al., 2017). This sparsity and diversity pose a challenge for studying interneuron subtypes in mammalian brain through electrode recordings. Here we demonstrate *in vivo* functional connectivity mapping between optically targeted interneuron pairs in neocortical layer 1 with QuasAr6a-based Optopatch. We also extended all-optical electrophysiology to the parvalbumin (PV)-expressing fast-spiking interneurons with QuasAr6b-based Optopatch. The faster kinetics of QuasAr6b enabled us to capture PV spike synchrony at sub-millisecond temporal resolution. Finally, we demonstrated optical detection of electrical synapses between PV neurons. Together, our study provides a suite of tools for all-optical interrogation of circuit dynamics in live mammalian brain.

## Results

### Photoselection enables video-based pooled high-throughput screens in mammalian cells

Previous attempts to engineer GEVIs have highlighted the importance of multi-dimensional screening (Herwig et al., 2017; Hochbaum et al., 2014; Kannan et al., 2018; McIsaac et al., 2014; Piatkevich et al., 2018; Villette et al., 2019) that incorporates static parameters (expression level, trafficking, brightness) as well as dynamic parameters (voltage sensitivity, response kinetics). Array-based screens, while readily compatible with real-time imaging, require laborious preparation of a large library of isolated genetic variants. In addition, screens for improved GEVIs are more likely to yield useful sensors for neurons if the variants are tested in mammalian cells instead of bacterial cells (Herwig et al., 2017; McIsaac et al., 2014).

Thus, we developed a video-based pooled screening platform for directed evolution of biosensors in mammalian cells. This platform consists of an ultra-widefield imaging component and a photoselection component (Figure 1A). The ultra-widefield imaging system, modified from the Firefly microscope described in (Werley et al., 2017b), enables characterization of the dynamic responses of many cells in parallel (>10^3^ cells per field of view) with millisecond time resolution.

**Figure 1.**
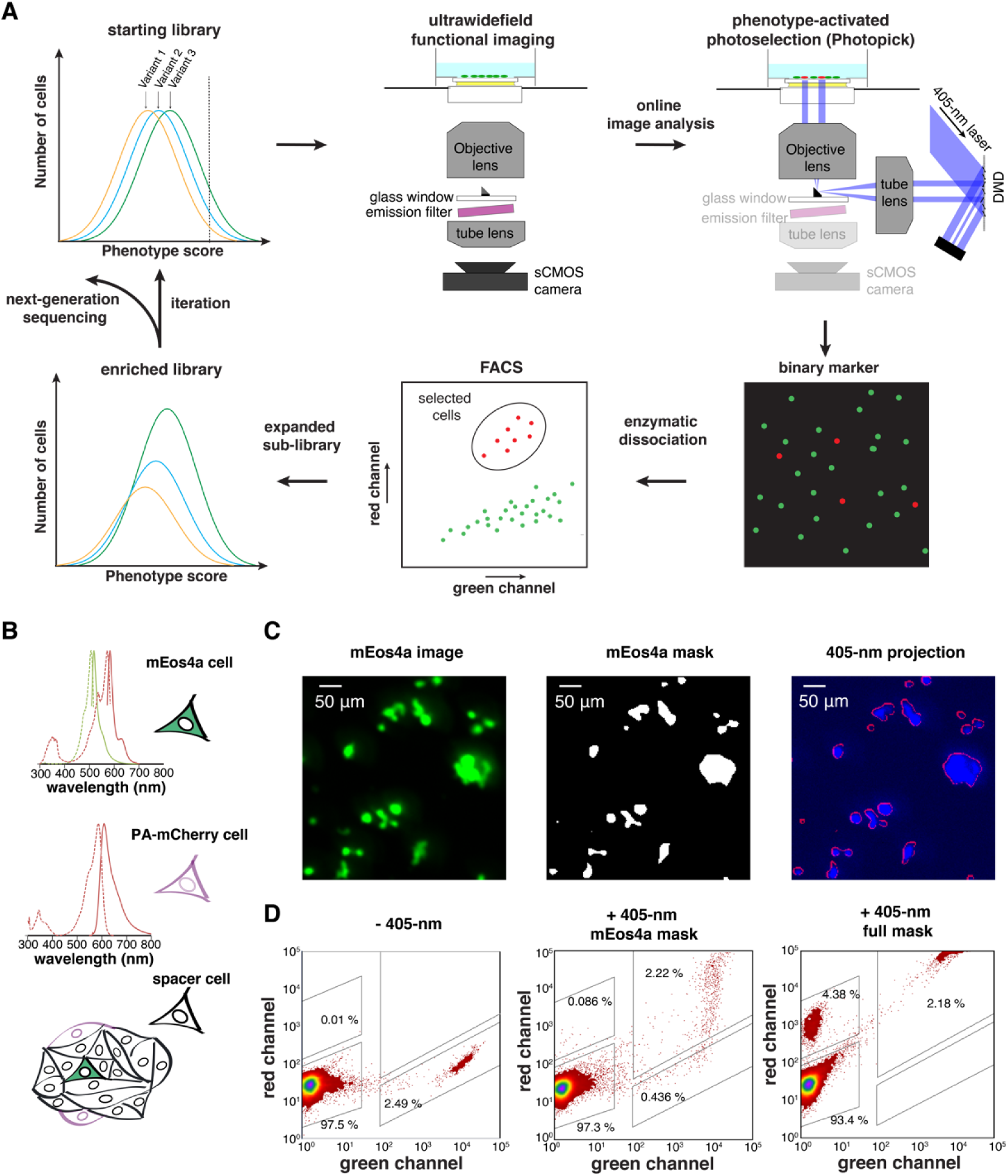
Photopick enables video-based pooled screening in mammalian cells. (A) Video-based pooled screening pipeline. (B) Test of optical targeting efficiency and fidelity using a mixed culture of cells expressing either of two phototaggable FPs, mEos4a or PA-mCherry. Spectra from FPbase (Lambert, 2019). (C) Selective phototagging of mEos4a^+^ cells in a hybrid monolayer of mEos4a^+^, PAmCherry^+^ and non-fluorescent HEK cells. (D) FACS analyses on the efficiency and fidelity of photoselection.

To isolate the desired phenotype from the pooled library, we developed “Photopick”, a phenotype-activated photoselection method. A phototaggable (photo-convertible, activable, or switchable) fluorescent protein (FP) is co-expressed with the mutant library in the host mammalian cells. Cells with the desired phenotype are selectively phototagged through patterned Illumination. Photopick creates a binary fluorescent marker to distinguish cells with desired phenotype. The phototagged cells are then sorted and recovered with fluorescence activated cell sorting (FACS). The phototagged population can be expanded to create a sub-library. In the course of culture expansion, the phototagged FP is degraded while fresh FP is synthesized, thus resetting the fluorescent marker to its initial state. This process can be iterated to further enrich for the desired phenotype. The shift of the prevalence distribution of candidate reporter genes is quantified with Illumina sequencing.

We used a digital micromirror array device (DMD) to address the target cells. We developed a protocol to achieve sub-pixel registration of the DMD and the wide-field imaging camera (average errors < 0.5 pixel, corresponding to 2.3 μm, Figure S1A), allowing precise optical targeting of single cells. We evaluated three candidate phototaggable FPs with different spectra for Photopick (Figure S1B): mEos4a, a green-to-red photoconvertible FP (Paez-Segala et al., 2015); PA-mCherry, a red photoactivable FP (Subach et al., 2009), and PA-GFP, a green photoactivable FP (Patterson et al., 2002). We found that mEos4a achieved efficient phototransformation at the lowest optical dose (fluence at 405 nm for 50% phototagging: 5.4 ± 0.3 J/cm^2^ for mEos4a; 18 ± 4 J/cm^2^ for PA-mCherry; 140 J/cm^2^ for PA-GFP, Figure S1C) while having a spectral window compatible with Arch-derived GEVIs.

We then evaluated the efficiency and fidelity of Photopick. We plated a mixture of mEos4a^+^ cells (green-to-red), PA-mCherry cells (dark-to-red) and blank HEK cells (approximate ratio 1:2:50) in a monolayer on the glass-bottomed dish (Figure 1B). We imaged the green (mEos4a) fluorescence and patterned illumination of violet light at the mEos4a^+^ pixels (Figure 1C). FACS analysis showed that the fidelity (phototagged mEos4a^+^ cells/(phototagged mEos4^+^ cells + phototagged PA-mCherry^+^ cells)) was approximately 96% and efficiency (phototagged mEos4a^+^ cells/all mEos4^+^ cells) was approximately 85% (Figure 1D). In a more stringent test, the blank HEK cells were omitted from the mixed monolayer (mEos4a^+^:PA-mCherry cells plated in a ratio 1:20). Targeted violet illumination of the mEos4^+^ cells again resulted in selective phototagging of mEos4a^+^ cells but not surrounding PA-mCherry^+^ cells (Figure S1D). We concluded that the Photopick system had the precision for phenotype-activated photoselection at cellular resolution.

### Directed evolution of Arch-derived GEVIs

High-throughput GEVI screening requires a means of inducing spikes in membrane potential. To achieve robust and reproducible electrical spikes in an easy-to-grow cell line, we adapted a synthetic electrophysiology model “spiking HEK” cell (Park et al., 2013; Zhang et al., 2016). Co-expression of a voltage-gated sodium channel Na_V_1.5 and an inward-rectifier potassium channel K_ir_2.1 enabled HEK cells to produce action potential-like spikes (Methods). We also stably expressed a blue-shifted channelrhodopsin, CheRiff, to optogenetically evoke the spikes. Whole-cell patch clamp (Figure 2B and Figure S2C, D) and voltage imaging with a red voltage-sensitive dye BeRST1 (Huang et al., 2015) (Figure 2C) validated the all-or-none spikes in response to increasing levels of optogenetic stimulation. Conveniently, the endogenously expressed gap junction proteins in HEK293 cells equalized changes of membrane potential in the connected cells (McNamara et al., 2018). Imaging of optogenetically triggered spiking with BeRST1 (Figure S2D) revealed a narrow distribution of fluorescence changes (Δ*F*/*F*_0_ = 0.25 ± 0.02, mean ± SD). Thus, CheRiff^+^ spiking HEK cells provided a uniform background for screening pooled GEVI libraries.

**Figure 2.**
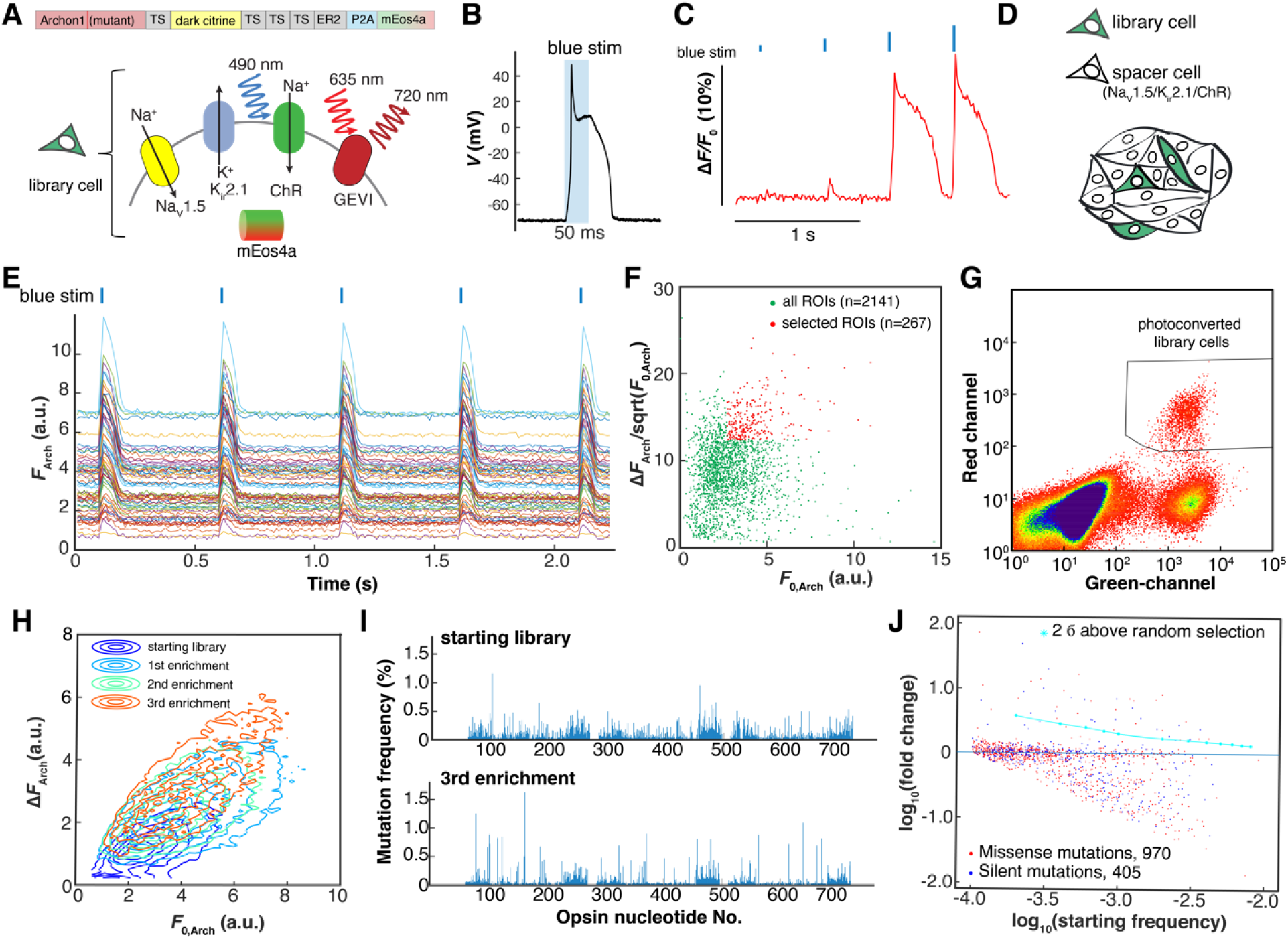
Directed evolution of an Archaerhodopsin-derived genetically encoded voltage indicator. (A) Genetic composition of the library cells. Spiking HEK cells containing a candidate GEVI mutant, channelrhodopsin actuator, mEos4a phototag, and Na_V_1.5 and K_ir_2.1 to imbue electrical excitability. Here TS is the Trafficking Sequence from K_ir_2.1(Hofherr et al., 2005), ER2 is the endoplasmic reticulum export motif from K_ir_2.0 (Gradinaru et al., 2010; Stockklausner et al., 2001), and P2A is a self-cleaving peptide. (B) Optogenetically triggered spike of the spiking HEK cell, recorded via whole-cell patch clamp (exc. at 488 nm, 4.4 mW/mm^2^, see also Figure S2C). (C) Threshold response of spiking HEK cells to 10-ms increasing optogenetic stimulus strengths, visualized with the voltage-sensitive dye BeRST1. (D) Sample preparation for pooled screening. Library cells were mixed with electrically excitable but non-fluorescent and optically inert spacer cells in approximately a 1:10 ratio. (E) Fluorescence traces extracted from individual sources in a pooled library screen. Δ*F*_Arch_ is calculated as the average baseline-to-peak difference in fluorescence and *F*_0, Arch_ is the average baseline fluorescence. (F) Scatter plot of Δ*F*_Arch_ /sqrt (*F*_0, Arch_) vs. *F*_0, Arch_ for all the ROIs in a 2.3-mm × 2.3-mm FOV. The quantity 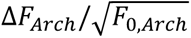 was used as a measure of shot-noise-limited SNR. (G) Representative FACS data showing three distinct population: photoconverted library cells; unselected library cells; spacer cells. (H) Three rounds of iterative enrichment shifted the population phenotype. (I) Manhattan plot showing the mutation frequency at each nucleotide. (J) Logarithmic plot of the starting mutation frequency vs. the fold of change. In this library, there were 970 missense mutations and 405 silent mutations.

As the starting template, we chose Archon1 (Piatkevich et al., 2018), which is an evolved version of Archaerhodopsin 3 that has been demonstrated for *in vivo* voltage imaging in mice (Fan et al., 2020; Piatkevich et al., 2019) and zebrafish larvae (Böhm et al., 2021). Since the voltage-sensing mechanism of rhodopsin-derived GEVIs is not fully understood (Maclaurin et al., 2013; Penzkofer et al., 2019, 2021), it is difficult to adopt a structure-guided approach to design the mutant library. In earlier screening efforts, beneficial mutations were found to scatter throughout the scaffold rather than to cluster at “hotspots” (Hochbaum et al., 2014; Piatkevich et al., 2018). Based on this observation, we introduced random mutations into the opsin sequence using error-prone PCR (Figure 2A, Figure S2A, B, Methods).

We developed a bicistronic vector to co-express single copies of GEVI mutants and mEos4a, and introduced these into spiking HEK cells via low-titer lentiviral transduction (multiplicity of infection ∼ 0.01; Figure 2A, Figure S2D, Methods). The expressing cells were separated from non-expressing cells by FACS and mixed with CheRiff^+^ spiking HEK cells lacking a GEVI (“spacer” cells) at a ratio of ∼1:10 (Figure 2D). The spacer cells served to homogenize the membrane potential across the monolayer and to create gaps between the library cells to facilitate image segmentation and photoselection.

The mutants were screened based on baseline brightness *F*_0_ and fractional fluorescence change, Δ*F*, induced by an optogenetically triggered action potential (Figure 2E, F). For each mutant, we calculated the quantity Δ*F*/√*F*_0_ as a measure of the shot noise-limited signal-to-noise ratio (SNR). From a single dish, 30,000 to 50,000 cells were scanned and approximately 12.5% were selected for photoconversion. The cells were then dissociated and sorted based on their fluorescence markers. Three distinct populations were observed in FACS (Figure 2G): the spacer cells (green^-^, red^-^), the unconverted library cells (green^+^, red^-^), and the photoconverted library cells (green^+^, red^+^). The converted library cells were recovered and then expanded for the next iteration of screening and selection. After three rounds of enrichment, we observed that the population had clearly shifted towards higher *F*_0_ and Δ*F* (Figure 2H). We sequenced the opsin mutations and analyzed the changes in single nucleotide polymorphism (SNP) prevalence (Figure 2I, J). While the majority of SNPs were either unaffected or depleted in the selection process, some missense mutations were positively selected above the 2σ threshold based on our stochastic stimulation (Methods).

### Engineering and Characterization of QuasAr6a and QuasAr6b in cultured cells

The sequencing results provided a short list of candidate mutations. We first created a panel (>30) of single missense mutants and screened them in HEK cells for expression and GEVI channel brightness. Next, we sought to combine these mutations. We found that combining two mutations that were close in linear sequence often resulted in detrimental effects. Therefore, we chose to combine distant mutations based on the Archaerhodopsin 3 structure (PDB 6GUY). We arrived at two new Archaerhodopsin-derived GEVIs (Figure 3A): QuasAr6a (Archon1 + W42G/M85I/- F98L/V124G/W148C/A238S) and QuasAr6b (Archon1 + W42G/M85I/F98L/V124G/W148C/ R237I). For subsequent characterization in cultured cells, we fused QuasAr6a and QuasAr6b with an optimized Citrine tag (Adam et al., 2019).

**Figure 3.**
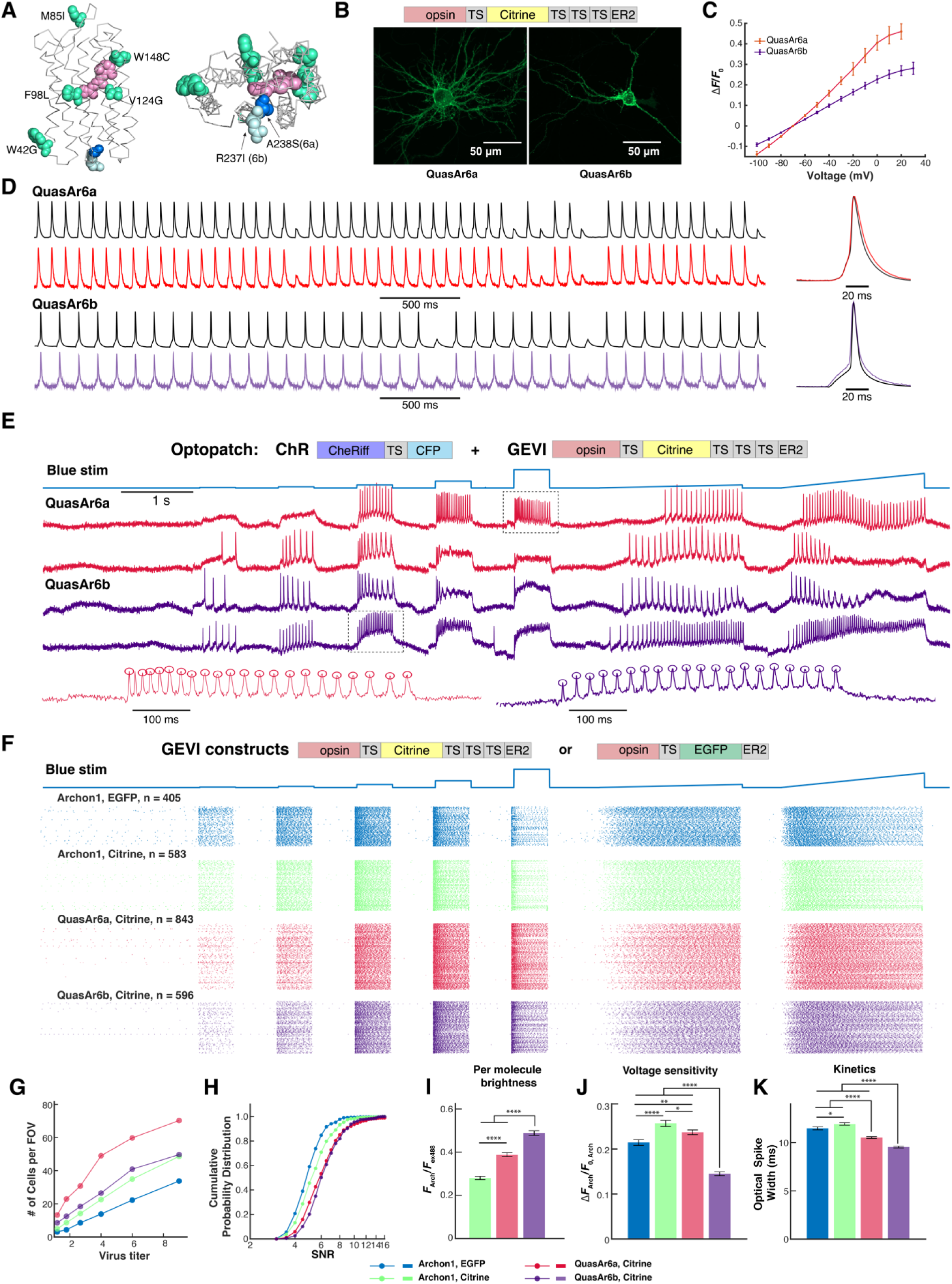
Characterization of QuasAr6a and QuasAr6b in cultured rat hippocampal neurons. (A) Homology model of QuasAr6a and QuasAr6b on the Archaerhodopsin 3 crystal structure (PDB: 6GUY), with the mutated residues for creating QuasAr6a and QuasAr6 from Archon1 highlighted. (B) Confocal image of Citrine fluorescence (z-projection) from QuasAr6a-Citrine and QuasAr6b-Citrine expressed in cultured rat hippocampal neurons. (C) Fluorescence vs. voltage curves for QuasAr6a (*n* = 3 cells) and QuasAr6b (*n* = 4 cells) measured under voltage clamp. Fluorescence is measured relative to *F*_0_ at the holding potential -70 mV. (D) Concurrent fluorescence and current-clamp recordings of a cultured hippocampal neuron expressing QuasAr6a-Citrine or QuasAr6b-Citrine in current-clamp mode. Left: Patch clamp (black) and fluorescence (red, purple) recordings. Right: overlay of spike-triggered average of the electrically and optically recorded spikes. Here spikes were induced by current injection (80 pA for 20 ms, 8.3 Hz). (E) Example traces from high-throughput Optopatch measurements using QuasAr6a or QuasAr6b combined with CheRiff-CFP in cultured rat hippocampal neurons. In the magnified views below, the circles indicate the automatically detected spike peaks. (F) Spike raster of the Optopatch measurements for five Archaerhodopsin-derived GEVIs at the highest titers tested. The GEVIs carried either TS-EGFP-ER2 or TS-Citrine-TS×3-ER2 tags. (G) Average number of cells with SNR > 3 per FOV with different GEVIs, as a function of virus titer. (H) Cumulative probability distribution of SNR for different GEVIs at the highest virus titer. (I-K) Comparison of per-molecule brightness, voltage sensitivity and optical spike widths at the highest lentivirus titer. *n* = 405 to 843 cells per condition. Error bars: SEM.

We performed biophysical characterization of QuasAr6a-Citrine and QuasAr6b-Citrine expressed in HEK cells (Figure S3). Both showed excellent membrane localization (Figure S3A). Compared with the template Archon1, QuasAr6a exhibited enhanced per molecule brightness (1.7-fold, Figure S3B), similar voltage sensitivity (73 ± 8% over 100 mV for QuasAr6a, *n* = 5 cells; 70 ± 13% over 100 mV for Archon1, *n* = 4 cells; mean ± SD, Figure 3C). QuasAr6b showed enhanced per molecule brightness (2.0-fold) and smaller fractional voltage sensitivity (24±4% over 100 mV) (Figure S3D), but faster on and off-kinetics (Figure S3D, E). Both QuasAr6a and QuasAr6b showed linear fluorescence-voltage (F-V) relations from -70 mV to 30 mV (Figure S3C) and excellent photostability (Figure S3F). Neither QuasAr6a nor QuasAr6b showed substantial blue light modulation of the near infrared fluorescence, in contrast to some other Arch variants (Adam et al., 2019; Chien et al., 2021; Pnevmatikakis et al., 2017) (Figure S3G). We also found no photocurrent under blue or red illumination for QuasAr6a or QuasAr6b (Figure S3H).

Next, we tested QuasAr6a-Citrine and QuasAr6b-Citrine in cultured rat hippocampal neurons. Both QuasAr6a and QuasAr6b showed efficient membrane localization (Figure 3B). QuasAr6a and QuasAr6b showed a fractional voltage sensitivity of 43 ± 4 % and 27 ± 3% from -70 mV to +20 mV (Figure 3C). Both QuasAr6a and QuasAr6b clearly reported single spikes and subthreshold events (Figure 3D).

To obtain robust statistics on the sensor performance, we used a high-throughput all-optical electrophysiology platform that could perform voltage imaging of > 100 cultured neurons in parallel at a 1 kHz frame rate (Werley et al., 2017a). We compared four Arch-derived GEVIs, Archon1-Citrine, QuasAr6a-Citrine, QuasAr6b-Citrine, and Archon1-EGFP that carried a TS-EGFP-ER2 tag as described in the original report (Piatkevich et al., 2018). Each of these GEVIs was paired with CheRiff via a bicistronic lentiviral expression vector to enable Optopatch measurement of neuronal excitability (Figure 3E, F). We tested each construct at 6 viral titers, 4 replicate wells for each condition. At the highest titer we measured for each construct between 405 and 843 cells, for a total of 2427 single-cell voltage imaging recordings.

Neurons were automatically segmented using an activity-based segmentation (Buchanan et al., 2018). Cells were kept if the single-trial, single-spike SNR (spike height:standard deviation of baseline noise) exceeded 3 (Methods). Since neurons in all wells were plated at the same density, the number of recorded cells per field of view (FOV) was an indicator of sensor performance. Across the titers, QuasAr6a consistently gave 2.1 - 4-fold more neurons with SNR above threshold per FOV (average 70 neurons per FOV at highest titer, *n* = 12 FOVs) compared to Archon1-EGFP (average 34 neurons per FOV at highest titer), and at least 1.5-fold more above-threshold neurons per FOV compared to Archon1-Citrine (average 49 neurons per FOV at highest titer). QuasAr6b also outperformed Archon1-EGFP by 1.5 - 2.2 fold (average 50 neurons per FOV at highest titer), and detected comparable above-threshold neurons per FOV as Archon1-Citrine. Among neurons detected with QuasAr6a-Citrine or QuasAr6b-Citrine, 6% or 3% of cells had an SNR between 3 and 4, respectively. By comparison, for Archon1-EGFP and Archon1-Citrine, 21% and 13% cells had an SNR between 3 and 4, respectively. Assuming Gaussian baseline noise, at an SNR of 3 and a 1 kHz frame rate, spike detection has a false positive rate of approximately 1 per 0.7 s, while at SNR of 4, the false positive rate falls to approximately 1 per 32 s. These data show that in cultured neurons both QuasAr6a and QuasAr6b were able to report the spike dynamics at higher fidelity than Archon1, while QuasAr6a also captured more spiking neurons.

We further analyzed the per molecule brightness (*F*_Arch_/*F*_ex488_), voltage sensitivity (Δ*F*_Arch_/*F*_0, Arch_), kinetics (optical spike width), baseline brightness, and expression level for the four sensors (Figure 3I-K, S4). QuasAr6a and QuasAr6b showed enhanced per-molecule brightness compared with Archon1 (1.4-fold and 1.7-fold, respectively; Figure 3I, S4B). For Archon1, substituting the EGFP tag with the Citrine tag improved voltage sensitivity by ∼20% across the titers and hence gave higher SNR (Figure 3J, S4C). The voltage sensitivity of QuasAr6a-Citrine was comparable to that of Archon1-Citrine and outperformed Archon1-EGFP at most titers (Figure 3J, S4C). The voltage sensitivity of QuasAr6b was lower than Archon1-Citrine or QuasAr6a-Citrine by 40∼45% (Figure 3J, S4C). Nonetheless, the superior brightness of QuasAr6b and greater expression level (1.46×) compensated for its lower voltage sensitivity to give a higher overall SNR. The optical spike width (Figure 3K, S4D) indicated that both QuasAr6a (optical spike width 10.4 ± 0.1 ms; mean ± SEM, *n* = 843 neurons) and QuasAr6b (9.5 ± 0.1 ms, *n* = 596 neurons) were faster than Archon1-Citrine (11.9 ± 0.1 ms, *n* = 583 neurons) or Archon1-EGFP (11.4 ± 0.1 ms, *n* = 583 neurons). Taken together, both QuasAr6a and QuasAr6b showed improved SNR and kinetics in cultured neurons compared to other Arch-based GEVIs.

### Validation of QuasAr6a and QuasAr6b *in vivo*

To resolve individual neurons in densely packed tissues, we designed soma-targeting versions for QuasAr6a and QuasAr6b by appending a Kv2.1 trafficking motif to the C-terminus (Adam et al., 2019; Baker et al., 2016; Lim et al., 2000; Piatkevich et al., 2019). somQuasAr6a and somQuasAr6b carried an EGFP tag as an expression marker. For optogenetic activation in tissue, we used a soma-localized version of CheRiff, somCheRiff (Adam et al., 2019). We made Cre-dependent bicistronic AAV constructs for Optopatch based on somQuasAr6a and somQuasAr6b, respectively. We sparsely expressed the Optopatch constructs in mouse cortex with AAV2/9 (Figure 4A, B). Confocal imaging of fixed and stained brain slices confirmed that somQuasAr6a/b and somCheRiff trafficked well *in vivo* and were restricted to soma and proximal dendrites.

**Figure 4.**
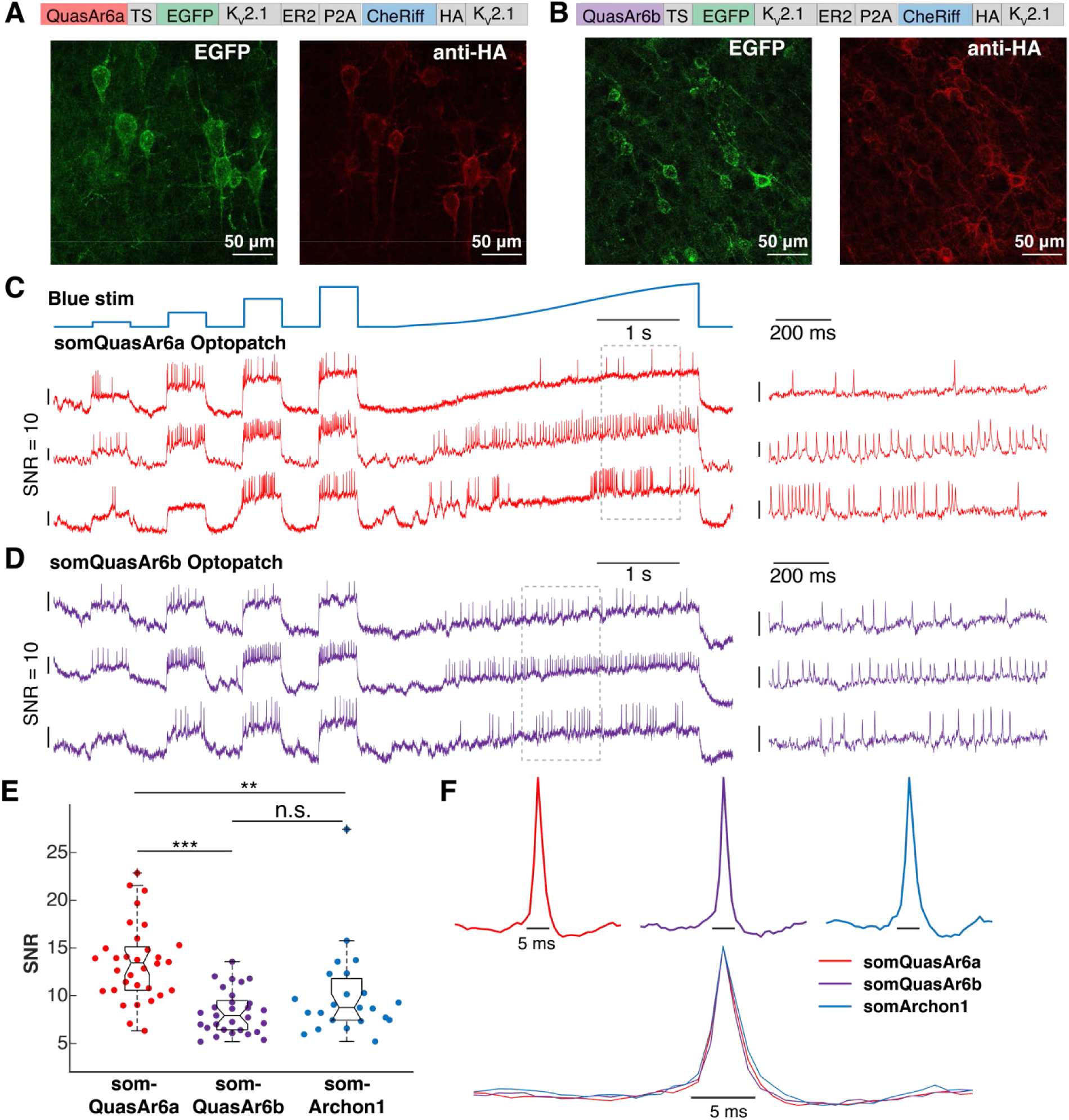
somQuasAr6a- and somQuasAr6b-based Optopatch in mouse cortex. (A, B) Bicistronic expression of soma-targeted QuasAr6a-EGFP (somQuasAr6a) or soma-targeted QuasAr6b-EGFP (somQuasAr6b) with somCheRiff-HA in cortical Layer 5. (C, D) Simultaneous optogenetic stimulation and voltage imaging in cortical L1 NDNF interneurons expressing somQuasAr6a- or somQuasAr6b-based Optopatch in anesthetized mice. Magnified view of the boxed regions is shown on the right. (E) Comparison of *in vivo* SNR of somQuasAr6a (*n* = 32 cells, 2 animals), somQuasAr6b (*n* = 29 cells, 2 animals), and somArchon1 (*n* = 23 cells, 2 animals) in NDNF interneurons. n.s. not significant; ** p < 0.01; *** p < 0.001 (Wilcoxon rank-sum test). (F) Spike-triggered average fluorescence waveforms of the optogenetically triggered spikes measured by somQuasAr6a, somQuasAr6b, and somArchon1.

Next, we sought to compare the *in vivo* performance of GEVIs in a genetically defined neuron population. We chose cortical neuron-derived neurotrophic factor (NDNF) cells for this purpose. In cortex, NDNF is a specific marker for GABAergic neurogliaform cells mostly restricted to the topmost 100 μm of layer 1 (L1) (Abs et al., 2018; Anastasiades et al., 2021; Schuman et al., 2019). The narrow distribution of depth made NDNF cells well suited for calibrating sensor performance. We expressed Optopatch constructs based on somQuasAr6a, somQuasAr6b, or somArchon1 in the visual cortex of NDNF-Cre^+/-^ mice. We used a holographic structured illumination system for *in vivo* voltage imaging and optogenetic stimulation (Figure S5A) (Fan et al., 2020). To ensure optimal focusing, cells were recorded one at a time, at a framerate of 1 kHz through a cortical window in head-fixed, anesthetized mice (Figure 4C, D). We calculated the SNRs for the optogenetically triggered spikes (Figure 4E). Optopatch constructs gave SNR values of: somQuasAr6a: 13.5 ± 4.0 (mean ± SD, n = 32 cells, 2 animals), somQuasAr6b: 8.3 ± 2.3 (mean ± SD, n = 29 cells, 2 animals) and somArchon1: 9.3 ± 2.8 (mean ± SD, n = 22 cells, 2 animals). In the samples expressing somQuasAr6b and somArchon1, many cells were near the analysis cutoff of SNR = 4, suggesting that the underlying distribution of SNR values for these reporters may have had lower mean than reported above. All three GEVIs reported spike waveforms with good fidelity (Figure S4F, S5B). These data suggest that for NDNF cells, the kinetics of all the three GEVIs are fast enough to faithfully report the waveform.

### Mapping functional connections between NDNF cell pairs with all-optical electrophysiology

The laterally projecting axon arbors of NDNF cells form a short-range, mutually inhibitory network within L1a on the outermost layer of the cortex. The role of this lateral inhibition in modulating the spike dynamics of L1 cells *in vivo* is not well understood. We previously showed that transient optogenetic activation of a population of L1 interneurons could suppress the spiking of a nearby cell (Fan et al., 2020). However, it was unclear how much effect activation of a single L1 interneuron would have on its neighbors.

Harnessing the improved SNR of somQuasAr6a in L1 NDNF cells, we mapped functional connections between optically targeted pairs of NDNF cells (Figure 5A). The postsynaptic cell was optogenetically stimulated with a steady step function of blue light to depolarize the cell and thereby increase the driving force for inhibitory chloride currents. The presynaptic cell was stimulated with a ramp waveform to evoke spiking when the blue light stimulus crossed an optical rheobase threshold. By inducing presynaptic spiking with a gradual blue light ramp, we could determine whether changes in spike rate of the postsynaptic cell coincided with the onset of presynaptic spiking. We monitored the voltage in both cells to observe whether there was a change in the spike-rate of the postsynaptic cell at the time of the first spike in the presynaptic cell. As control measurements, we included epochs without presynaptic stimulation. We then swapped the blue light waveforms between the pair to test connectivity in the other direction (Figure 4B). For each pair of cells, we performed 2-7 trials. We used a Bayesian Adaptive Kernel Smoother (BAKS) (Ahmadi et al., 2018) to estimate the instantaneous spike rate. We performed these measurements in anesthetized mice to minimize the fluctuations from other synaptic or neuromodulatory inputs.

**Figure 5.**
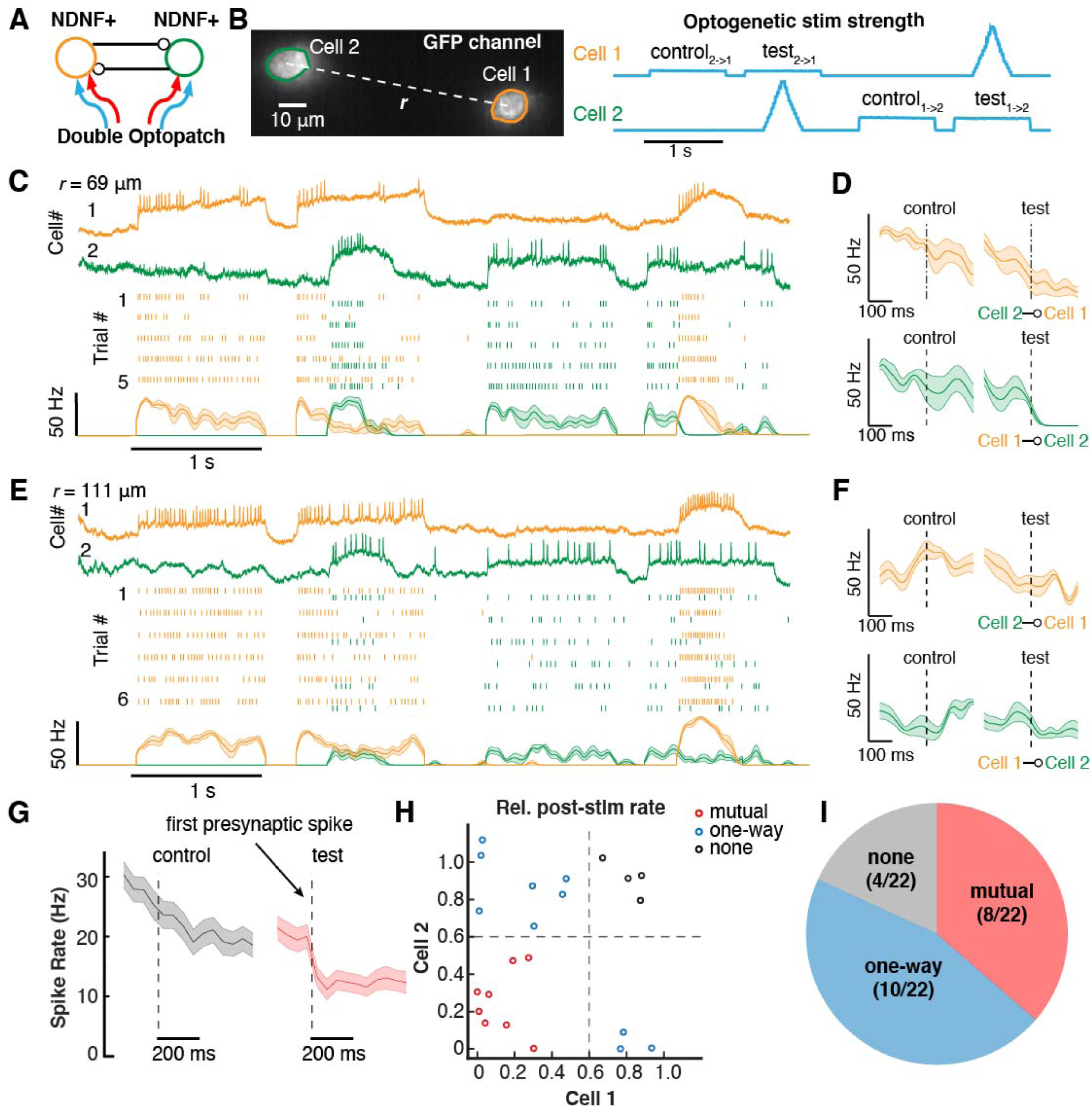
Optical dissection of inhibitory connections between NDNF interneurons. (A) Scheme of two-cell Optopatch in NDNF interneurons. (B) Optogenetic stimulation waveform for probing inhibitory connections in a pair of optically targeted NDNF cells. The presynaptic cell was activated by ramp stimulation. The postsynaptic cell was depolarized with a constant step stimulation to increase the driving force for inhibitory currents. (C) Optopatch revealed strong mutual inhibition between a pair of NDNF neurons (intersoma distance *r* = 69 μm). Top to bottom: representative traces, raster plots for 5 consecutive trials; spike rate estimated by Bayesian Adaptive Kernel Smoother (shading: SEM). (D) For the cell pair in (C), change of spike rate after the onset of the first presynaptic spike (dashed line) or at the corresponding time in the control epoch. (E, F) Similar to (C, D), for a pair of NDNF cells with weaker inhibitory connections (inter-soma distance *r* = 111 μm). (G) Average of all the postsynaptic cells (n = 51 cells). (H) Inhibition strength for 22 pairs of cells. The inhibition strength was quantified by the relative post-stimulation rate, defined as the ratio of the minimal spike rate within the window 0 to 100 ms after the first presynaptic spike, to the average spike rate during the window -100 ms to -10 ms before the first presynaptic spike. (I) Pie chart showing the distribution of the mutual, one-way and undetectable inhibition between 22 cell pairs.

We performed double Optopatch experiments on 30 pairs of cells from 4 animals, where the inter-soma distance ranged from 46 μm to 216 μm. Figure 5C, D shows a pair with strong reciprocal inhibition, while Figure 5E, F shows a pair exhibiting one-way inhibition. In order to accurately test for a synaptically driven decrease in spike rate, we restricted analysis to postsynaptic cells in which optogenetic stimulation evoked a spike rate above 5 Hz (n = 51 cells, 22 pairs where both cells spiked above 5 Hz). Activation of a putative presynaptic cell reduced the mean spike rate of its neighbor from 20 ± 2 Hz to 11 ± 1.7 Hz (mean ± SEM). We found that 36% of the pairs (8/22) showed approximately symmetrical mutual inhibition, 45% of all the pairs (10/22) showed one-way inhibition, and the rest showed no inhibition (4/22). Thus, while NDNF cells on average strongly inhibited neighboring NDNF neurons, this inhibitory effect was not always reciprocal. These statistics of functional connectivity will be important in constraining models of L1 dynamics.

### Probing hippocampal PV neuron dynamics all-optical electrophysiology

PV fast-spiking neurons play a key role in regulating the excitation/inhibition balance of the brain through inhibition of excitatory pyramidal neurons (Ferguson et al., 2018). While two photon-targeted whole-cell patch clamp recording has been demonstrated in live mouse cortex (Jouhanneau et al., 2018)*, in vivo* electrophysiological recordings from the subcortical regions have been rarely reported due to the sparsity of PV cells and the anatomical constraints for two photon-targeted patch clamp. Voltage imaging of hippocampal PV neurons has been performed with the one-photon GEVI Voltron-JF525 (Abdelfattah et al., 2019), Voltron-JF552 (Abdelfattah et al., 2021), or the two-photon GEVI ASAP3 (Villette et al., 2019). However, for neither ASAP3 nor Voltron-could voltage imaging be paired with optogenetic stimulation. Moreover, due to PV neuron’s narrow spikes (FWHM < 0.5 ms) (Antonoudiou et al., 2020) and the relatively slow kinetics of ASAP3, voltage imaging of PV cells with ASAP3 gave a low SNR, potentially reducing the fidelity of spike detection.

We explored whether the fast variant QuasAr6b could enable accurate detection of voltage dynamics in hippocampal PV neurons. We injected AAV2/9 for Cre-dependent somQuasAr6b-Optopatch and somArchon1-Optopatch into the hippocampus of a PV-Cre^+/-^ transgenic mouse. The cortical tissue above the hippocampus CA1 region was carefully removed and replaced with a cannula window (Adam et al., 2019; Dombeck et al., 2010). We performed voltage imaging at 2 kHz framerate for PV^+^ neurons expressing the Optopatch constructs in Stratum Oriens (Figure 6A). Optogenetic stimulation readily triggered high-frequency spikes as visualized by somQuasAr6b (Figure 6B). We compared the SNR and optical width of optogenetically triggered spikes detected by somQuasAr6b (n = 24 cells, 3 animals) and somArchon1 (n = 23 cells, 2 animals). somQuasAr6b showed higher SNR (8.0 ± 2.5; mean ± SD) and narrower optical FWHM (0.91 ± 0.15 ms, mean ± SD) than Archon1 (SNR, 5.4 ± 1.5; optical FWHM, 1.1 ± 0.1 ms, Figure 6C, D). The average optogenetically triggered spike rate of PV neurons followed the intensity of blue light (Figure 6E). We also imaged the optogenetically triggered spikes with somQuasAr6b at a framerate of 4 kHz (n = 13 cells, 2 animals, Figure S6). We found that the mean spike half-width reduced from 0.91 ms at 2 kHz to 0.71 ms at 4 kHz (Figure S6B). This optical spike width approaches the spike width of hippocampal PV neurons measured by patch clamp (0.49 ± 0.04 ms) (Antonoudiou et al., 2020). Thus, somQuasAr6b is a fast and high-SNR sensor particularly suited for reporting sub-millisecond voltage dynamics.

**Figure 6.**
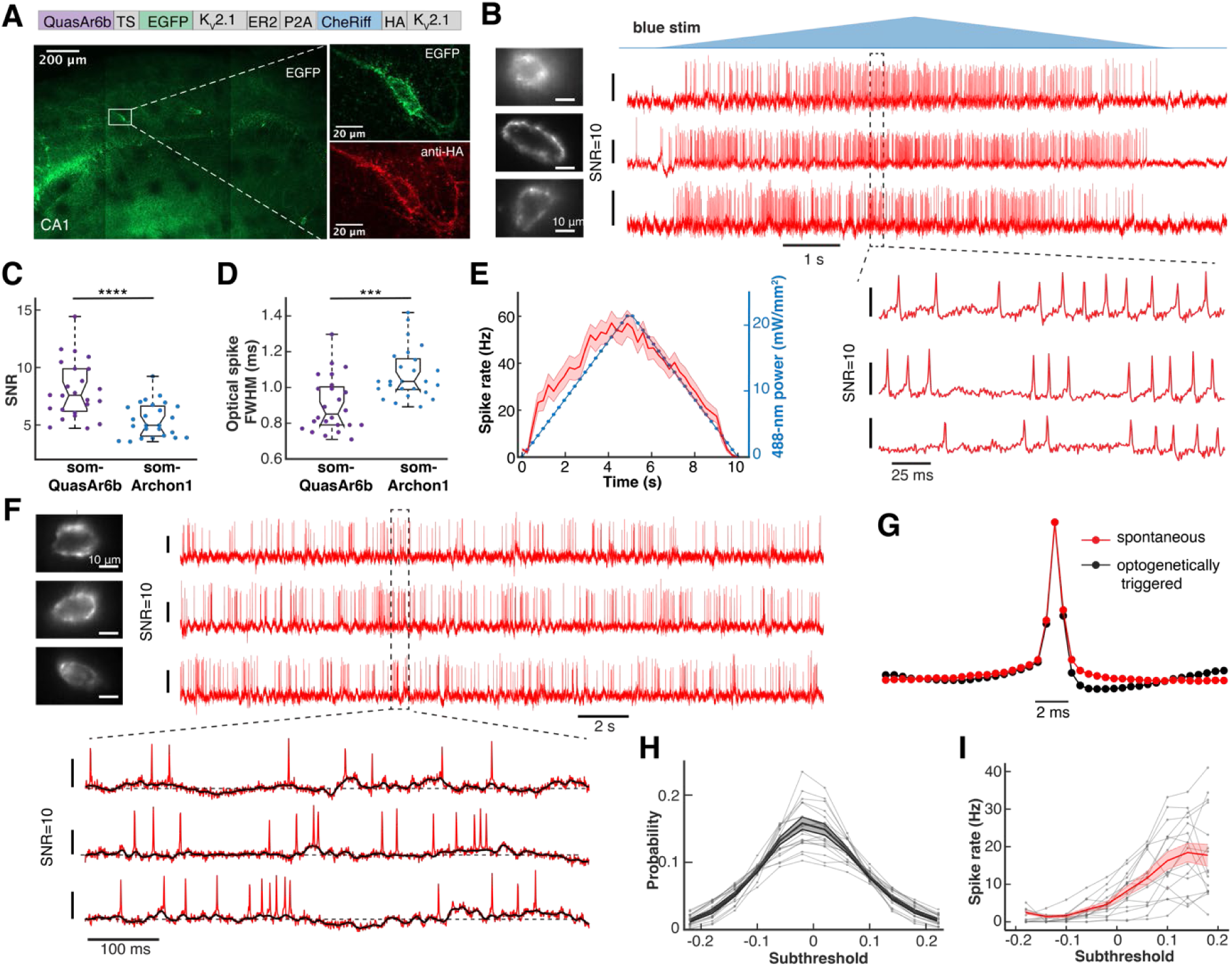
Optogenetically triggered and spontaneous activity in hippocampal CA1 PV neurons, recorded at 1973 Hz. (A) Sparse expression of somQuasAr6b-based Optopatch in hippocampus CA1 oriens of PV-Cre mice. (B) Double-ramp Optopatch measurements in PV cells in anesthetized mice. (C) Comparison of the *in vivo* SNR of QuasAr6b (*n* = 24 cells, 3 animals) and Archon1 in hippocampal PV cells (*n* = 25 cells, 2 animals). **** p < 0.0001 (Wilcoxon rank-sum test). (D) Comparison of the *in vivo* kinetics of QuasAr6b and Archon1 in hippocampal PV cells. *** p < 0.001 (Wilcoxon rank-sum test). (E) Average optogenetically triggered spike rate of as a function of blue light intensity in PV neurons expressing somQuasAr6b-based Optopatch (*n* = 25 cells, 2 animals). Shading: SEM. (F) Voltage imaging of spontaneous voltage dynamics of PV neurons in awake mice. In the magnified view, the black line indicates the subthreshold dynamics. (G) spike-triggered average fluorescence waveform of spontaneous spikes in awake mice or optogenetically triggered spikes in anesthetized mice. (H) Histogram of subthreshold levels during spontaneous activity in awake mice (*n* = 16 cells, 3 animals). Shading: SEM. (I) Individual (grey) and average (red) dependence of spike rate on optically inferred sub-threshold voltage (*n* = 16 cells, 3 animals). Shading: SEM. In (H-I) the subthreshold voltage is measured relative to the mean spike amplitude.

We next recorded the spontaneous voltage dynamics of PV cells (n = 16 cells, 3 animals) in awake mice with somQuasAr6b (Figure 6F). We found that the spontaneous and optogenetically triggered spikes showed similar spike width. However, the optogenetically triggered spikes showed more pronounced after-hyperpolarization (Figure 6G). Unlike the bimodal up/down states observed for cortical PV cells (Zucca et al., 2017), the subthreshold potential of hippocampal Oriens PV cells was approximately Gaussian distributed (Figure 6H). The PV spike rate increased as a sigmoidal function of subthreshold potential (Figure 6I). These results demonstrate that QuasAr6b could report PV cell electrical dynamics in awake mice.

### Hippocampal PV cells showed a high degree of spike synchrony

Voltage imaging provided a unique opportunity to compare the precise spike timing between multiple simultaneously recorded PV cells. In a rate-based spiking model, one would expect that at a low average spike rate, spiking “coincidences”, i.e., two or more cells spiking at almost the same time, should be extremely rare. On the other hand, if precise spike timing mattered, one might find reliable events in which multiple cells spiked in precise temporal sequences, or even at the same time.

Figure 7 shows an example pairwise recording of hippocampal PV cells in awake mice with 10 synchronous spikes (spike maxima within 0.5 ms) and 97 nearly synchronous spikes (spike maxima within 5 ms) in a 60 s recording where the two cells spiked at average rates of 5.6 Hz and 3.0 Hz, respectively. Among PV cell pairs separated by < 340 μm, on average 21.0 ± 8.5% of spikes were nearly synchronous, whereas after randomly shuffling the interspike intervals only 3.9 ± 2.4% of spikes were nearly synchronous (5 ms window, mean ± SD; *n* = 45 cell pairs, 2 animals; mean spike rate 4.6 ± 2.7 Hz, mean ± SD; Figure S7A, Methods). For the six most synchronized pairs (37.9 ± 2.1% of spikes nearly synchronous, mean ± SEM), time-shifting either spike train by only 8 ms decreased the fraction of nearly synchronous spikes to 16.8 ± 2.8%, establishing that spike synchrony arose from precise spike-to-spike correlations between cells, as opposed to global fluctuations in mean spike rate (Figure S7B).

**Figure 7.**
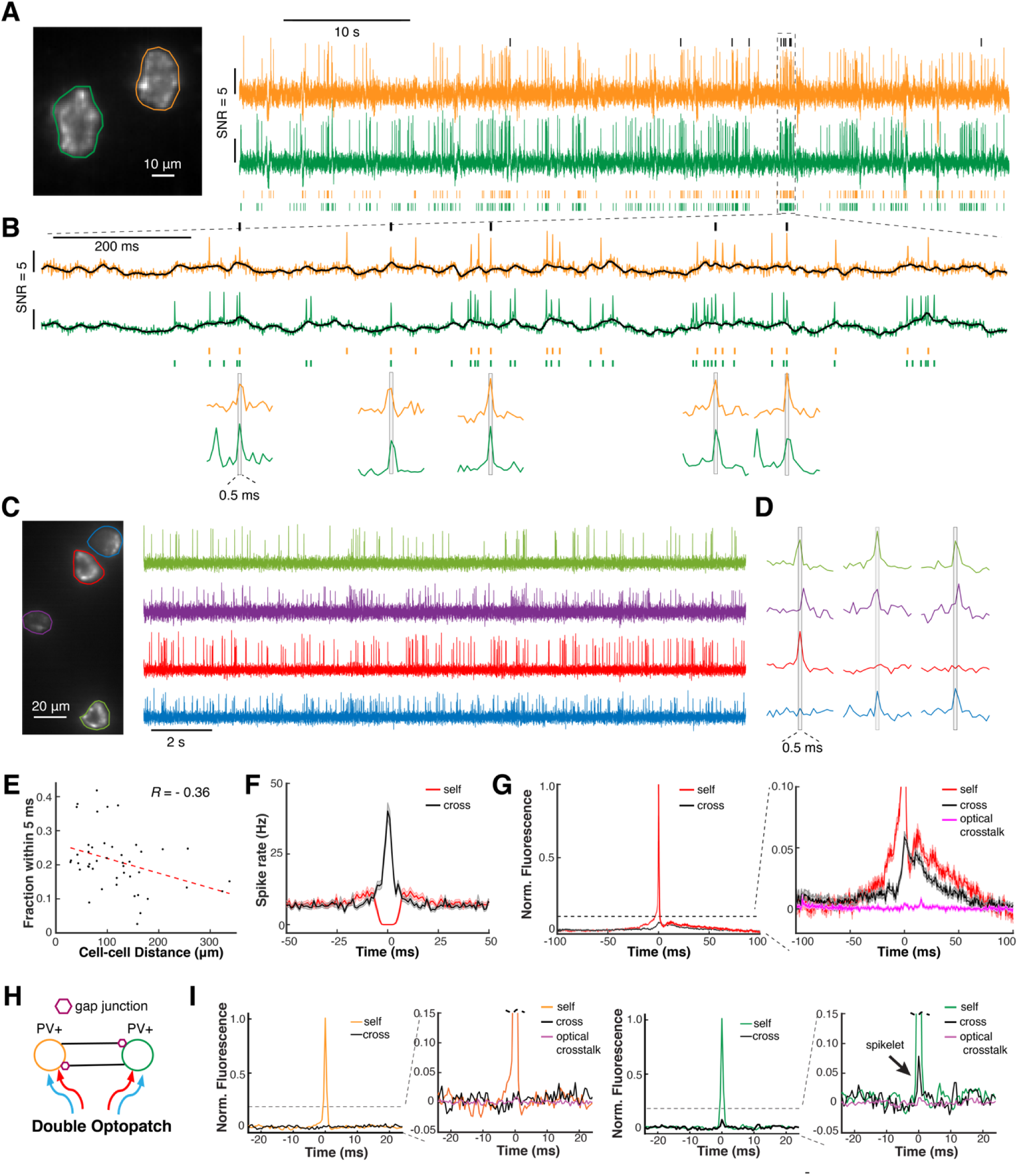
Correlated voltage dynamics of two or more hippocampal PV cells. (A) Simultaneous recording of spontaneous dynamics in two PV neurons in a lightly anesthetized mouse. Black ticks indicate synchronous spikes (< 0.5 ms time-difference in peaks). (B) Magnified view of the boxed region in (A). (C) Simultaneous recording from four hippocampal PV cells. (D) Magnified view of synchronous spikes in trios of PV cells. (E) Fraction of nearly synchronous spikes (< 5 ms time-difference in peaks) vs. the inter-soma distance (*n* = 45 pairs, 2 animals). (F) Self- and cross- spike-triggered spike rate for PV cell pairs (bin = 1 ms; *n* = 45 pairs, 2 animals). (G) Self- and cross spike-triggered voltage waveform for events where the spike maxima in each cell were separated by > 10 ms (*n* = 31 pairs from 2 animals). (H) Double Optopatch experiment to probe gap junction connections between PV cells. (I) In a pair of PV cells 44 μm apart, a gap-junction-mediated spikelet was observed in one cell but not the other.

Figure 7C shows an example recording from four PV cells. Among the six two-cell subsets, 22 ± 3% (mean ± SD) of the spikes were nearly synchronous. Remarkably, among the four trios, 7.4 ± 2.0% (mean ± SD) of the spikes were nearly synchronous. There were three 4-way near-synchronous spikes among an average of 132 spikes/cell in the 20 s recording. Time-shifting the spike trains of all possible three-cell subsets by 5 ms, 10 ms, and 15 ms, respectively, decreased the fraction of three-way near-synchronous spikes to 3.2 ± 0.9 % and the number of 4-way near-synchronous spikes to zero (Figure S7C, Methods). These results suggest that collective “network spikes” may be an important mode of PV cell activity.

As the intercellular separation increased, we observed only a slight decrease in the magnitude of the sub-threshold cross-correlations (Figure S7D) and also only a slight decrease in the proportion of pairwise synchronous spikes (Figure 7E). These observations indicate correlated inputs to and outputs of the PV network over length scales of at least 340 μm.

Several non-exclusive factors could contribute to near-synchronous network spikes: PV cells might have shared synaptic inputs whose positive fluctuations drive near-synchronous spikes; gap junction coupling between PV cells could further smooth cell-to-cell variations in subthreshold potentials, enhancing spike synchrony; and gap junction couplings could cause a spike in one cell to induce spikelets in neighboring cells, increasing their likelihood of near-synchronous spiking (Hjorth et al., 2009; van Welie et al., 2016). To explore the origin of the network spikes, we studied the joint statistics of subthreshold voltages and spike times in simultaneously recorded pairs of cells. First, we calculated the mean firing rate at time t + *τ* in cell *j*, given a spike at time *t* in cell *i*, i.e., the self (*i* = *j*) and cross (*i* ≠ *j*) spike-triggered firing rate (STFR, Figure 7F). The self-STFR showed the expected refractory period followed by a decay toward the average spike rate of 6.4 ± 6.5 Hz (*n* = 60 cells, 2 animals, mean ± SD, SEM = 0.9 Hz). The cross-STFR showed a sharp peak to 40 Hz at *τ* = 0, with a full-width at half-maximum of 4.5 ms, consistent with the high rate of near-synchronous spikes.

To explore the sub-threshold dynamics driving these near-synchronous spikes, we then calculated the mean self- and cross spike-triggered voltage waveform (STVW). We separately calculated these functions for events where only one cell spiked (Figure 7G, spike maxima in the two cells separated by > 10 ms) and for events with spike maxima in the two cells separated by < 10 ms (Figure S7E). For the cross-STVW when only Cell 1 spiked, the Cell 2 (“cross”) voltage showed little depolarization for *τ* < 0, had a sharp jump near *τ* = 0, and then relaxed closely following the waveform of Cell 1 for *τ* > 0 (Figure 7G). For the cross STVW when both cells spiked, the sub-threshold waveforms preceding (*τ* < 0) and following (*τ* > 0) the spike closely tracked for both cells (Figure S7E). To test whether the subthreshold dynamics in Cell 2 were influence by scattered fluorescence from Cell 1, we calculated the cross-STVW in a region midway between the two cells (Figure S7F). The signal from the intervening mask was negligible (Figure 7G), consistent with prior results (Fan et al., 2020).

The appearance of a low-pass-filtered copy of the Cell 1 waveform in Cell 2, even when Cell 2 did not spike, is strongly suggestive of a gap junction-mediated coupling mechanism. In contrast to the measurements on the NDNF cells, we did not observe signs of inhibitory synaptic coupling between the PV cells.

### Visualizing gap junction-mediated spikelets in PV cells

Without knowing the correlational structure of the synaptic inputs, it was not possible to unambiguously establish the role of direct electrical coupling between the PV cells. To address this uncertainty, we turned to targeted optogenetic stimulation of individual PV cells. In Optopatch recordings on pairs of PV cells in lightly anesthetized animals (Figure 7H, I, Figure S7G-I), we used optogenetic stimulation to trigger spikes alternately in each cell and recorded the spiking and subthreshold dynamics in both. Consistent with the measures of spontaneous activity, we did not observe suppression of spontaneous spike rate through lateral inhibition (Figure S7G).

We calculated the self- and cross-STVW (Figure 7I, S7H, I) for eight PV pairs. Five pairs had an inter-soma distance below 100 μm, and three pairs were separated by more than 100 μm (119-377 μm). For three pairs separated by less than 100 μm, we observed a spikelet in one of the two cells, consistent with dual patch clamp recordings of gap junction-coupled neurons (van Welie et al., 2016) (Figure 7I, S7H). The amplitude of the spikelets was 3.6 to 6.4% (mean 4.6%) of the full spike amplitude in the post-synaptic cell. The waveform of the spikelets resembled that of the optogenetically triggered spikes, and the timing of the spikelets corresponded to the timing of the other cell’s spikes at millisecond precision, consistent with gap junctional coupling. These results suggest that gap junctional coupling was indeed present among the hippocampal PV cells, and might mediate their collective network spikes.

## Discussion

We engineered two complementary Arch-derived GEVIs using a video-based, pooled screening platform (“Photopick”) for mammalian cells. Pooled screens offer the practical advantage of lower cost and higher throughput compared to arrayed screens (Feldman et al., 2019; Hasle et al., 2020; Kanfer et al., 2021; Lawson et al., 2021; Lee et al., 2020; Piatkevich et al., 2018; Yan et al., 2021). Similar methodology could be used to optimize fluorescent sensors of other modalities. To achieve spectral compatibility with blue or green reporters, one could mark target cells with a dark-to-green (e.g., PA-GFP) or dark-to-red (e.g., PA-mCherry) FP instead of the green-to-red mEos4a FP we used here. The Photopick platform is in principle compatible with any imaging-based assay of cellular structure or dynamics and a variety of downstream assays, e.g, single-cell sequencing. Thus, potential applications include forward genetic screens, e.g., for genes that affect cell migration, chemotaxis or responses to mechanical or metabolic perturbations.

In any recording, it is important to consider whether the unrecorded samples may contribute to systematic errors. The improved performance of QuasAr6a and QuasAr6b significantly increased the number of cultured neurons which produced sufficient SNR for a recording, and thereby reduced the sampling bias of voltage imaging. We calibrated the *in vivo* performance of GEVIs in two types of interneurons with distinct electrophysiological properties. In the fast-spiking PV neurons with narrow spikes, the faster sensor QuasAr6b, despite of its lower voltage sensitivity, exhibited better SNR. Our data highlights the interdependence of SNR and sensor kinetics as well as the importance of cell type-specific characterization for future development of voltage imaging. To our surprise the mutations selected in our screen were all located on the protein surface, rather than close to the chromophore. This insight may prove helpful for the ongoing GEVI engineering efforts.

Functional connectivity mapping in mammalian brain has remained a longstanding challenge in neuroscience. With a few exceptions (Ferrarese et al., 2018; Jouhanneau et al., 2018; Jouhanneau et al., 2019), most monosynaptic mapping work has been performed in acute slice. Yet, connectivity in acute slices may be underestimated due to severed connections, and the functional significance of observed connections can only be assessed *in vivo*. We observed strong spike rate suppression (∼45%) in neighboring cells by a single active NDNF cell in anesthetized mice. We also found that this inhibitory effect are heterogenous and sometimes asymmetrical between pairs of NDNF cells, consistent with results in acute slice (Schuman et al., 2019). A key next step will be to relate the functional connectivity maps to the responses of the network to naturalistic sensory and modulatory inputs.

Electrical synapses comprised of gap junctions are widespread between interneurons, including between cortical and hippocampal PV cells (Fukuda et al., 2000; Fukuda et al., 2006,Shigematsu, 2019 #114). Gap junctions have been shown by patch clamp measurements to promote spike and subthreshold synchronization *in vivo* (Hjorth et al., 2009; Tong et al., 2020; van Welie et al., 2016 ) in a distance-dependent manner. However, functional studies of gap junction in live brain have been limited by the difficulty of paired whole-cell recordings. All-optical electrophysiology enabled us to confirm the electric coupling between hippocampal PV cells, which may account for the remarkable abundance of spontaneous millisecond-precision spike synchrony. The existence of collective “network spikes” in gap junction-coupled PV cells is a phenomenon that can only be detected *in vivo*. The influence of these network spikes on CA1 pyramidal cells and ultimately on hippocampal dynamics remains to be determined.

## Acknowledgments

We thank B.L. Sabatini and O. Yizhar for advice and discussion; C. Dulac for the PV-Cre mouse line; M. Andermann and J. Fernando for advice on cranial window surgery; T.D. Green, K. Williams, Urs Böhm, A. Preecha, H. Dahche, Y. Adam, and G. Testa-Silva for technical assistance and advice; M.P. Chien for advice on pooled screening; E. Miller for the BeRST1 dye; B. Arnold from Harvard FAS Informatics for assistance with Illumina sequencing data analysis; the Bauer Core Facility at Harvard University for FACS service; the Biopolymers Facility at Harvard Medical School for next-generation sequencing; Harvard Center for Biological Imaging (RRID:SCR_018673) for infrastructure and support on confocal imaging.

## Author contributions

H.T. and A.E.C. conceived and designed the study. H.T. designed and conducted all the experiments except for the high-throughput Optopatch assay in cultured neurons. B.G. and V.P. assisted with the optics on the ultra-widefield microscope for pooled screening. H.C.D, H.T., and D.W.C improved the structured illumination microscope for *in vivo* imaging based on an earlier version built by L.Z.F. H.C.D. developed the Matlab control software for the structured illumination microscope. H.T., C.A.W., and G. B. B. designed the high-throughput Optopatch assay in cultured neurons for characterizing GEVIs. H. U, H. S., and J.J. performed the high-throughput Optopatch assay. Y.Q. assisted with the *in vivo* imaging experiments. P. P. performed electrophysiological experiments in acute brain slice. S.B. prepared the cultured neurons for GEVI characterization and performed the mouse husbandry. L.Z.F and K.D. contributed to the *in vivo* validation of the GEVIs in the early stage. H.T. and A.E.C. analyzed the data and wrote the manuscript. A.E.C. supervised the research.

## Data and materials availability

Plasmids encoding QuasAr6a or QuasAr6b will be available on Addgene upon publication. Data, code, and materials used in the analysis are available upon reasonable request to A.E.C.

## Supplementary Materials

Materials and Methods Figs. S1 to S7

---

**Figure S1.**
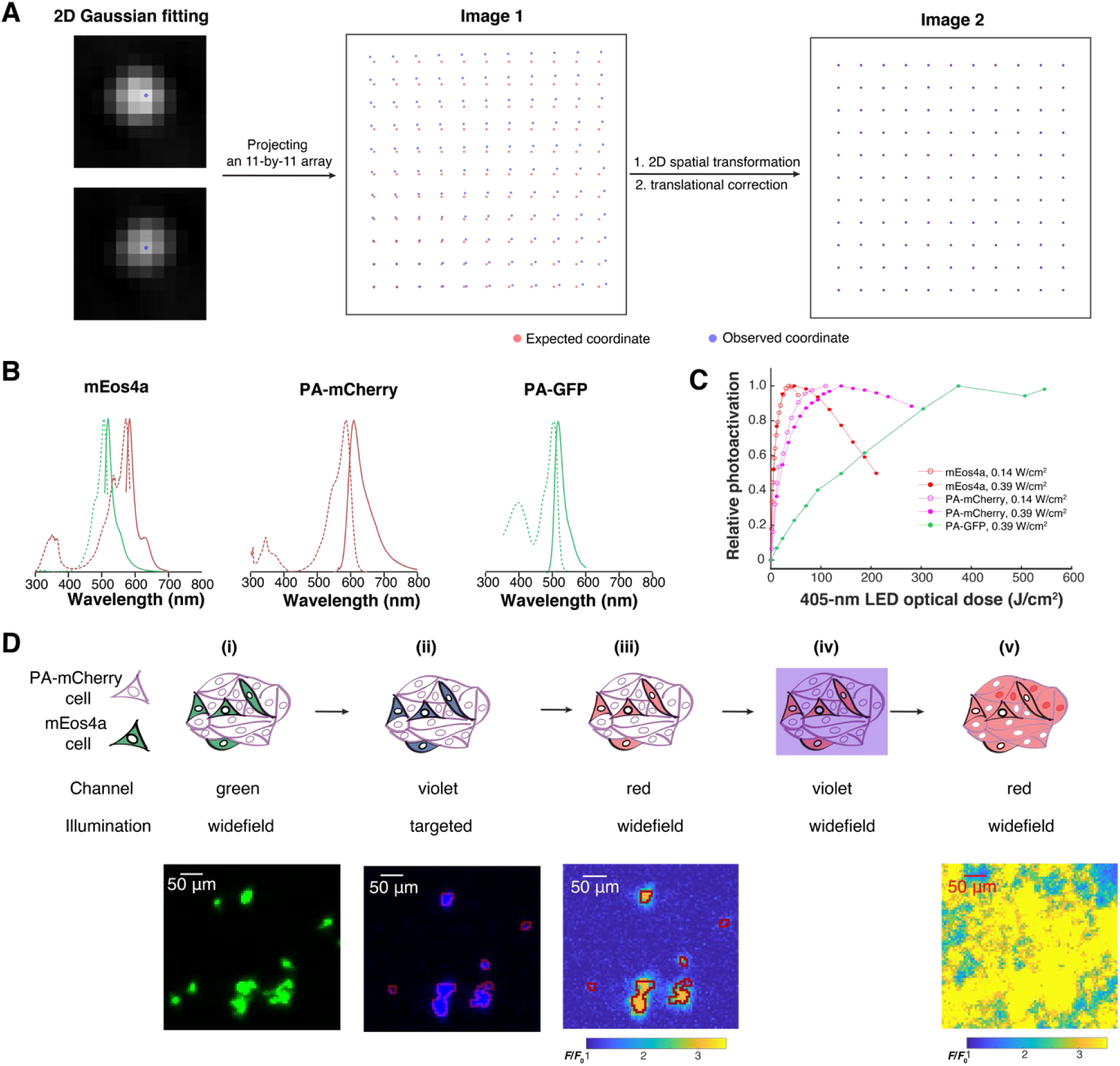
Calibration of Photopick, an imaging-based method for isolating mammalian cells from pooled culture, related to Figure 1. (A) Procedure for registering the DMD and camera pixels. An 11 x 11 grid of spots was projected onto a homogeneous exposure target. The observed locations in the camera were used to develop a piecewise-linear transformation to map DMD pixels onto camera pixels. In this example, the registration reduced the average projection error from 11.6 pixels to 0.22 pixels. (B) Fluorescence excitation and emission spectra of three phototaggable FPs, PA-GFP, PA- mCherry, and mEos4a. For mEos4a, the spectra are given in the pre-activation state (green) and post-activation state (red). For the other FPs, the activated spectra are shown. (C) Phototransformation efficiency vs. optical dose of 405-nm LED light. (D) Selective phototagging of mEos4a^+^ cells embedded in PA-mCherry^+^ cells (mEos4a^+^ : PA- mCherry^+^ = 1:20). Based on the green channel image (i), a mEOS4a mask was created for targeted photoconversion of mEos4a with violet (ii). The red channel image shows that the phototagging was highly specific (iii). The monolayer of cells was then broadly illuminated with violet light (iv) to drive the photoactivation of PA-mCherry^+^ cells (v).

**Figure S2.**
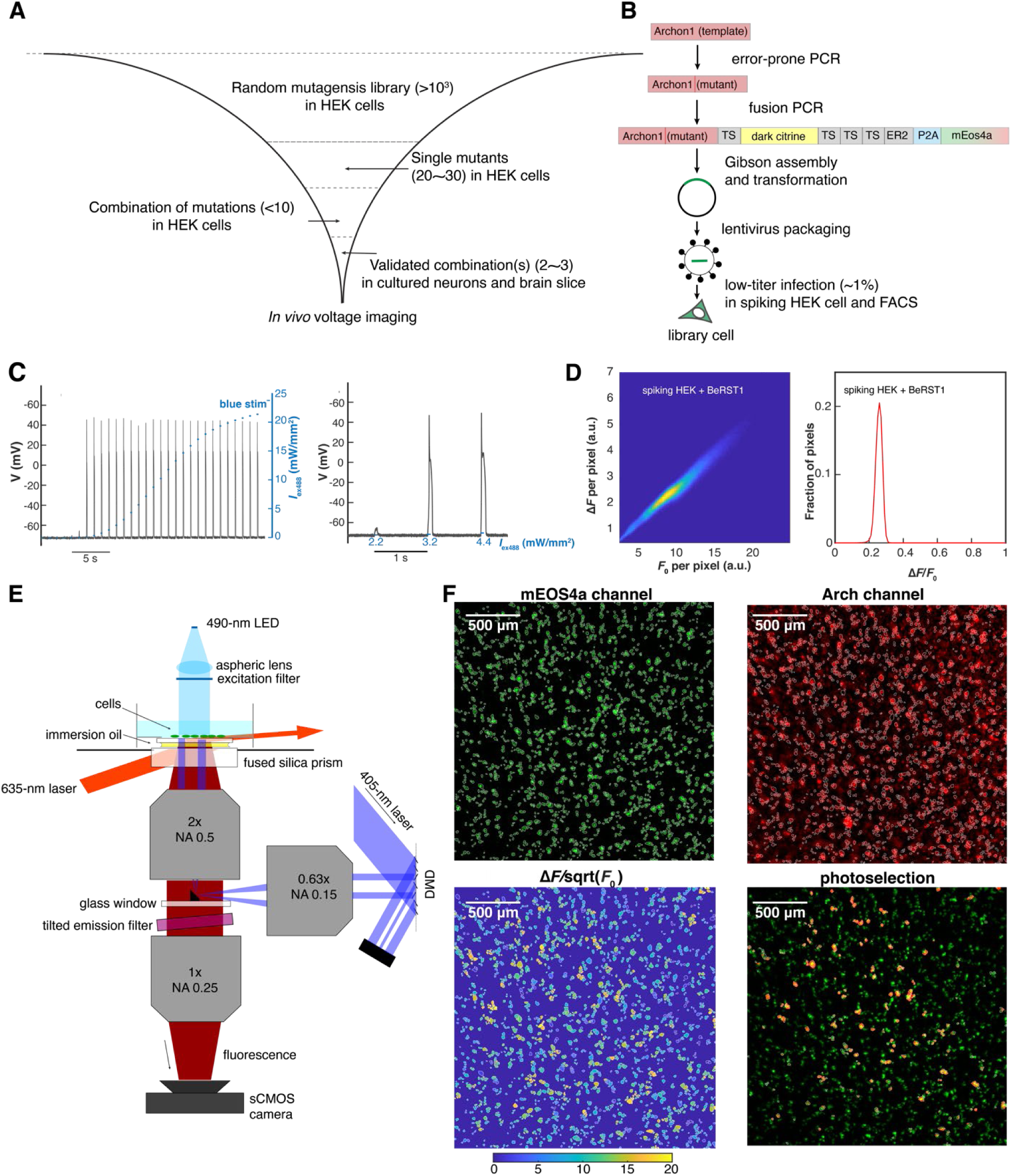
Video-based pooled screening of Arch-derived GEVIs, related to Figure 2. (A) Work-flow of engineering improved archaerhodopsin-derived GEVIs. (B) Generation of the library cells. (C) Current clamp measurement of membrane potential in spiking HEK cells reveals “all-or-none” spiking in response to increasing optogenetic stimulation (488 nm). Left: membrane potential change measured for the optical dose curve (0 - 22 mW/mm^2^). Right: enlarged view showing the threshold transition. (D) Distribution of membrane potential changes in a spiking HEK cell monolayer, reported with a voltage-sensitive dye BeRST1. BeRST1 fluorescence was recorded in the same configuration as for screening. Left: heatmap of Δ*F* vs. *F*_0_ for all pixels in a 2.3×2.3 mm FOV (500×500-pixels). Right: histogram of Δ*F*/*F*_0_. (Mean ± SD: 0.25 ± 0.02). (E) Optical system for all-optical pooled screening. (F) Image analysis for phenotype-activated photoselection. Top left: ROIs generated by image segmentation in the mEos4a channel (exc: 490 nm). Top right: baseline fluorescence (*F*_0_) image in the Arch channel (exc: 635 nm). Bottom left: heatmap of Δ*F*/sqrt(*F*_0_) for individual ROIs. Here Δ*F*/sqrt(*F*_0_) is used as a proxy for shot noise-limited for SNR. Bottom right: overlay of the patterned violet light (pseudocolor red) and mEos4a image.

**Figure S3.**
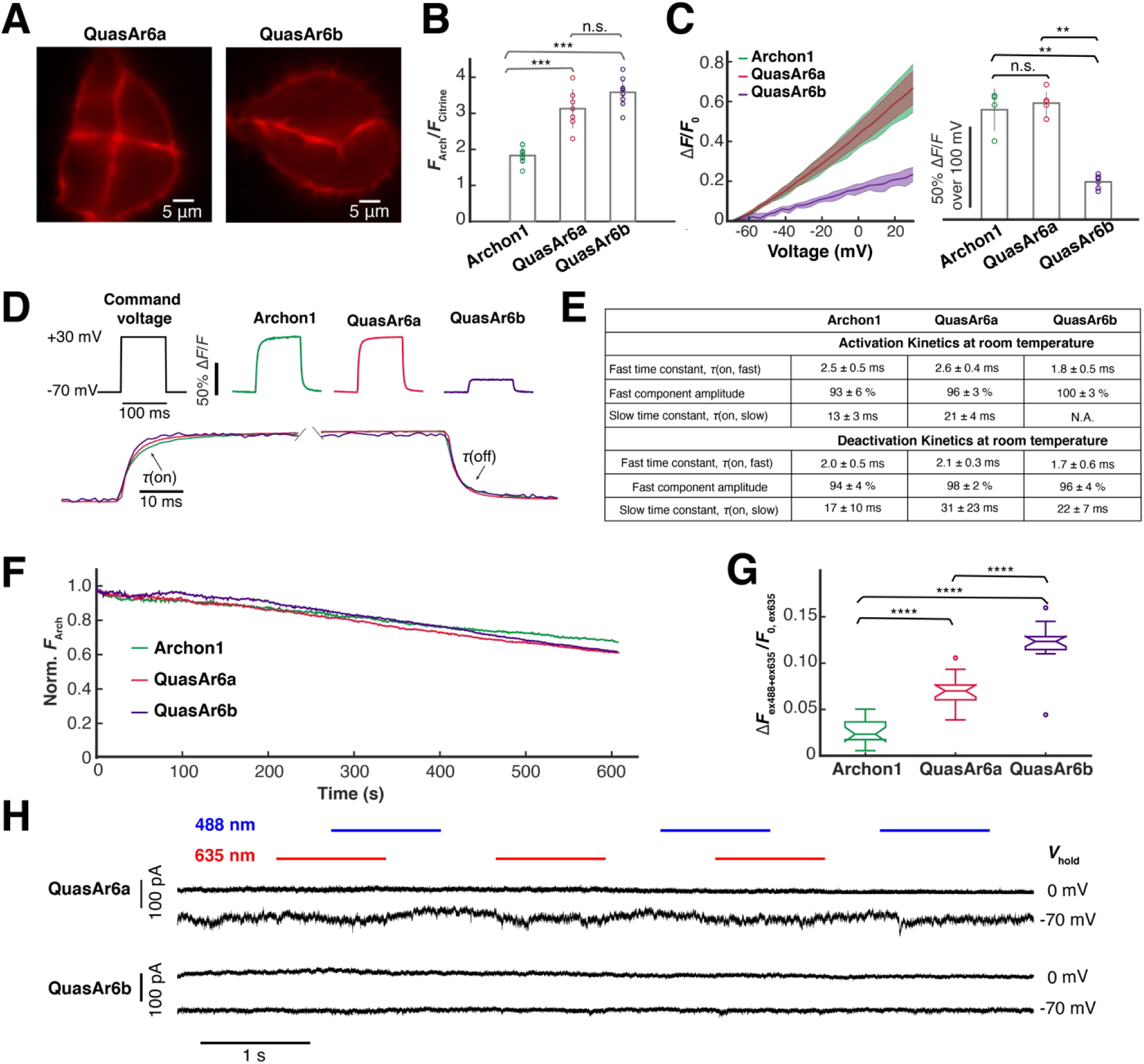
Characterization of QuasAr6a-Citrine and QuasAr6b-Citrine in HEK293T cells, related to Figure 3. (A) Arch-channel (exc: 635 nm, em: 670 – 746 nm) fluorescence images of QuasAr6a-Citrine and QuasAr6b-Citrine expressed in HEK cells. (B) Relative brightness per molecule of Archon1-Citrine (*n* = 10 cells), QuasAr6a-Citrine (*n* = 7 cells), and QuasAr6b-Citrine (*n* = 10 cells) measured as a ratio of whole-cell *F*_Arch_ to *F*_Citrine_. n.s. not significant, p > 0.05; *** p < 0.001 (Wilcoxon rank-sum test). (C) Voltage sensitivity measured by concurrent voltage clamp and fluorescence. Left: Fractional fluorescence change vs. membrane voltage; shading: SD. Right: Voltage sensitivity (ΔF/F per 100 mV: Archon1-Citrine, *n* = 4 cells; QuasAr6a-Citrine, *n* = 5 cells; QuasAr6b-Citrine, *n* = 6 cells). n.s. not significant, p > 0.05; ** p < 0.01 (Wilcoxon rank-sum test). (D) Voltage step-response kinetics measured by recording the fluorescence change during a 100 ms voltage step from -70 mV to +30 mV. Top: fluorescence. Bottom: overlay of fluorescence traces. (Archon1-Citrine, *n* = 4 cells; QuasAr6a-Citrine, *n* = 5 cells; QuasAr6b-Citrine, *n* = 6 cells). Measurements were performed at room temperature. (E) Summary of the step-response kinetic data, fitted with a biexponential model. (F) Photobleaching by 635 nm laser (420 W/cm^2^) over 10 min (*n* = 2 cells for each construct). All constructs showed < 40% photobleaching over 10 min with 635 nm illumination (420 W/cm^2^). (G) Quantification of blue-light induced photoactivation (635 nm, 420 W/cm^2^; 488 nm, 0.37 W/cm^2^). The blue-activation coefficient is defined as the Arch-channel fluorescence signal change under both blue and red excitation (*F*_ex488+ex635_ - *F*_ex488_ - *F*_ex635_), normalized by the baseline Arch-channel fluorescence (*F*_0, ex635_) (Chien et al., 2021). Blue-activation coefficient (mean ± SD): 0.05 ± 0.01 for Archon1 (*n* = 15 cells); 0.07 ± 0.02 for QuasAr6a (*n* = 15 cells); 0.12 ± 0.02 for QuasAr6b (*n* = 16 cells). ****p < 0.0001 (Wilcoxon rank-sum test). (H) HEK cells expressing QuasAr6a or QuasAr6b showed no photocurrents under either 488 nm, 635 nm or combined illumination at either -70 mV or 0 mV holding potentials (488 nm: 124 W/cm^2^; 635 nm: 1500 W/cm^2^).

**Figure S4.**
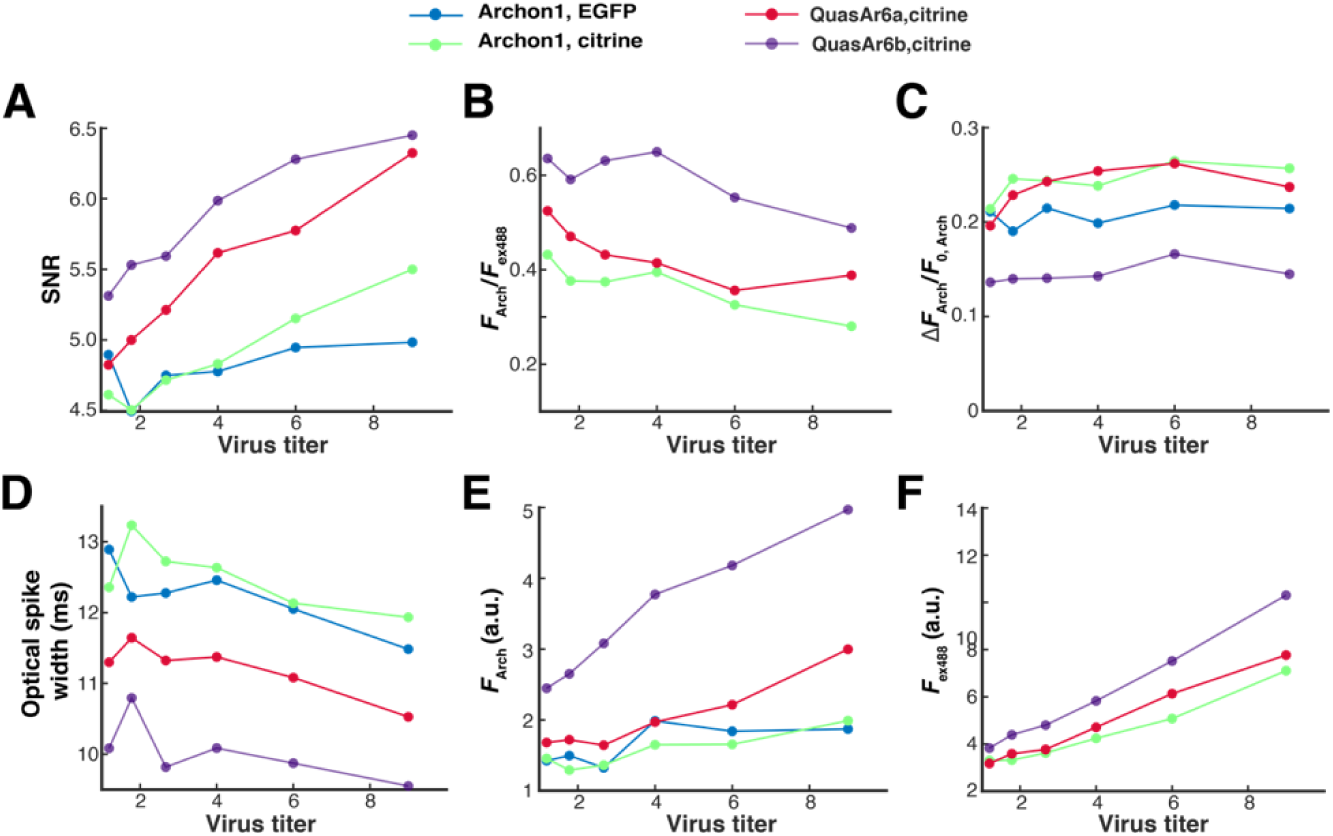
Metrics of GEVI performances in the high-throughput Optopatch assay, related to Figure 3. Relative titers: 1.19, 1.78, 2.67, 4, 6, 9. Statistics (Archon-EGFP, Archon-Citrine, QuasAr6a-Citrine, QuasAr6b-Citrine): *n* = 37, 59, 159, 103 cells (relative titer = 1.19); *n* = 52, 104, 275, 150 cells (relative titer = 1.78); *n* = 103, 170, 369, 221 (relative titer = 2.67); *n* = 166, 271, 588, 318 (relative titer = 4); *n* = 267, 419, 717, 491 (relative titer = 6); *n* = 405, 583, 843, 596 (relative titer = 9). (A) SNR: spike height divided by the root mean square (RMS) baseline noise. (B) *F*_Arch_/*F*_ex488_: per-molecule brightness as a ratio of baseline fluorescence in the Arch channel to the baseline fluorescence in the Citrine channel. (C) Δ*F*_Arch_/*F*_0, Arch_: voltage sensitivity as a ratio of the increase in fluorescence during a spike to the baseline fluorescence. (D) Optical spike width: full width measured at 80% below the action potential peak. (E) *F*_0, Arch_: baseline fluorescence in the Arch channel (exc: 635 nm). (F) *F*_ex488_: baseline fluorescence in the Citrine channel (exc: 488 nm).

**Figure S5.**
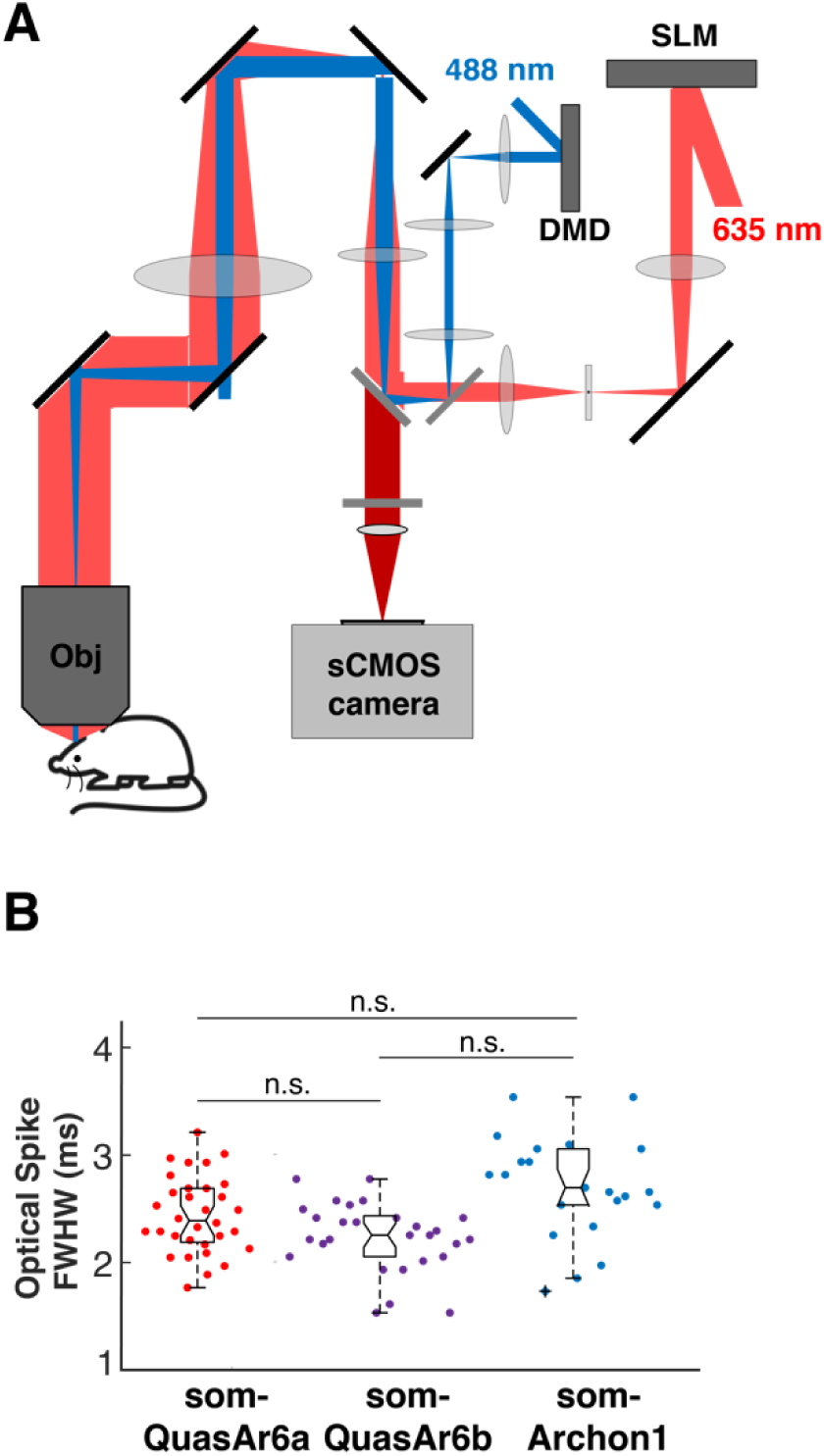
somQuasAr6a- and somQuasAr6b-based Optopatch in mouse cortex, related to Figure 4. (A) Optical system for patterned optogenetic stimulation and holographic voltage imaging. (B) Optical spike full-width at half-maximum (FWHM) of optogenetically triggered spikes in NDNF cells, imaged with somQuasAr6a (*n* = 32 cells, 2 animals), somQuasAr6b (*n* = 29 cells, 2 animals), and somArchon1 (*n* = 23 cells, 2 animals) at a 1 kHz frame rate. n.s. not significant (Wilcoxon rank-sum test).

**Figure S6.**
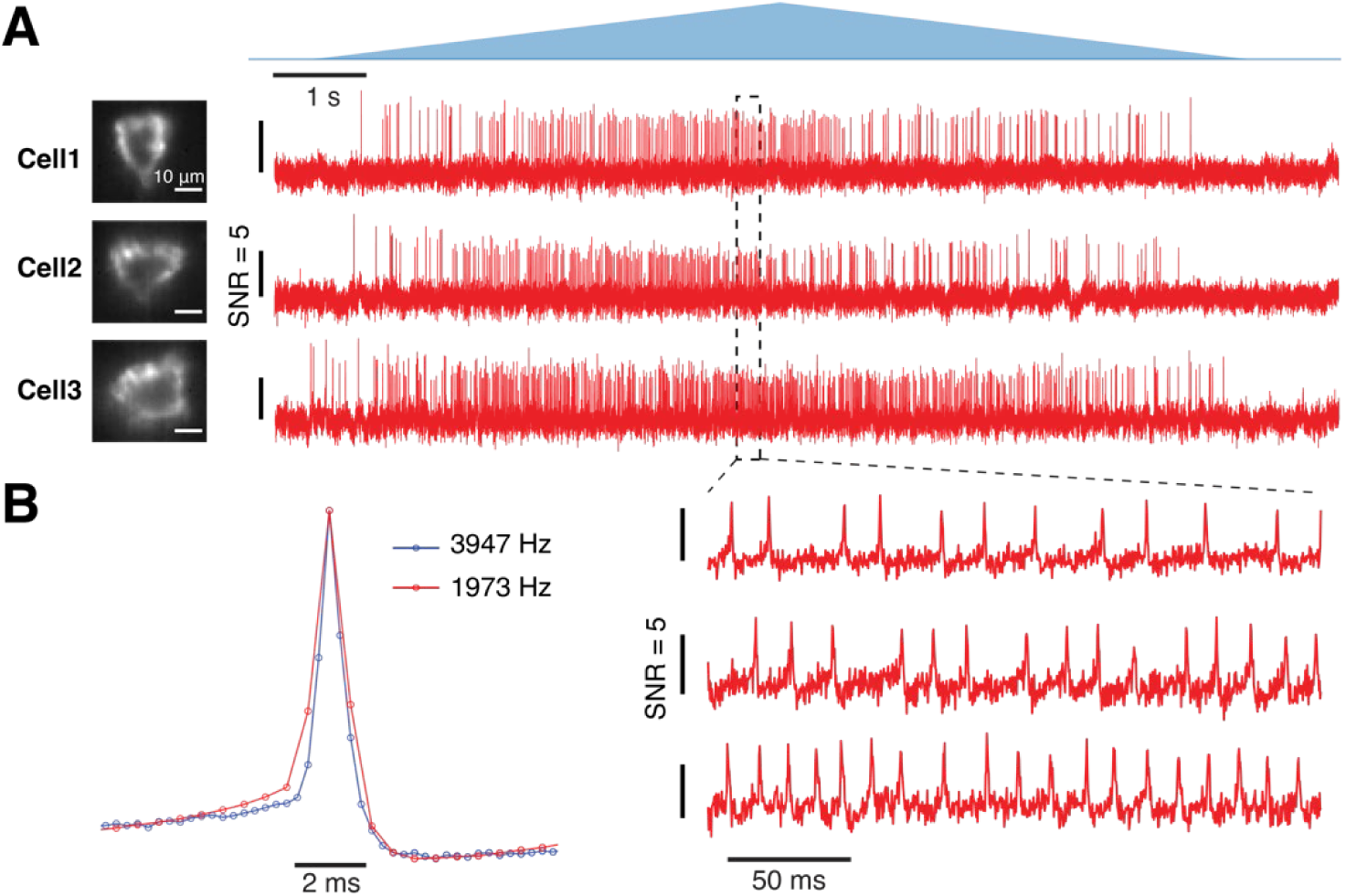
Optopatch in hippocampal PV cells at 4 kHz, related to Figure 6. To fit the entire cell body into the 4 kHz (3947 Hz) recording zone of the sCMOS camera, a 10x, NA 0.6 objective was used. For the 2 kHz (1973 Hz) recordings, a 25x, NA 1.05 objective was used. (A) Representative Optopatch traces. (B) Spike-triggered average fluorescence waveform of optogenetically trigged spikes recorded at 3947 (*n* = 13 cells, 2 animals) or 1973 Hz (*n* = 24 cells, 3 animals).

**Figure S7.**
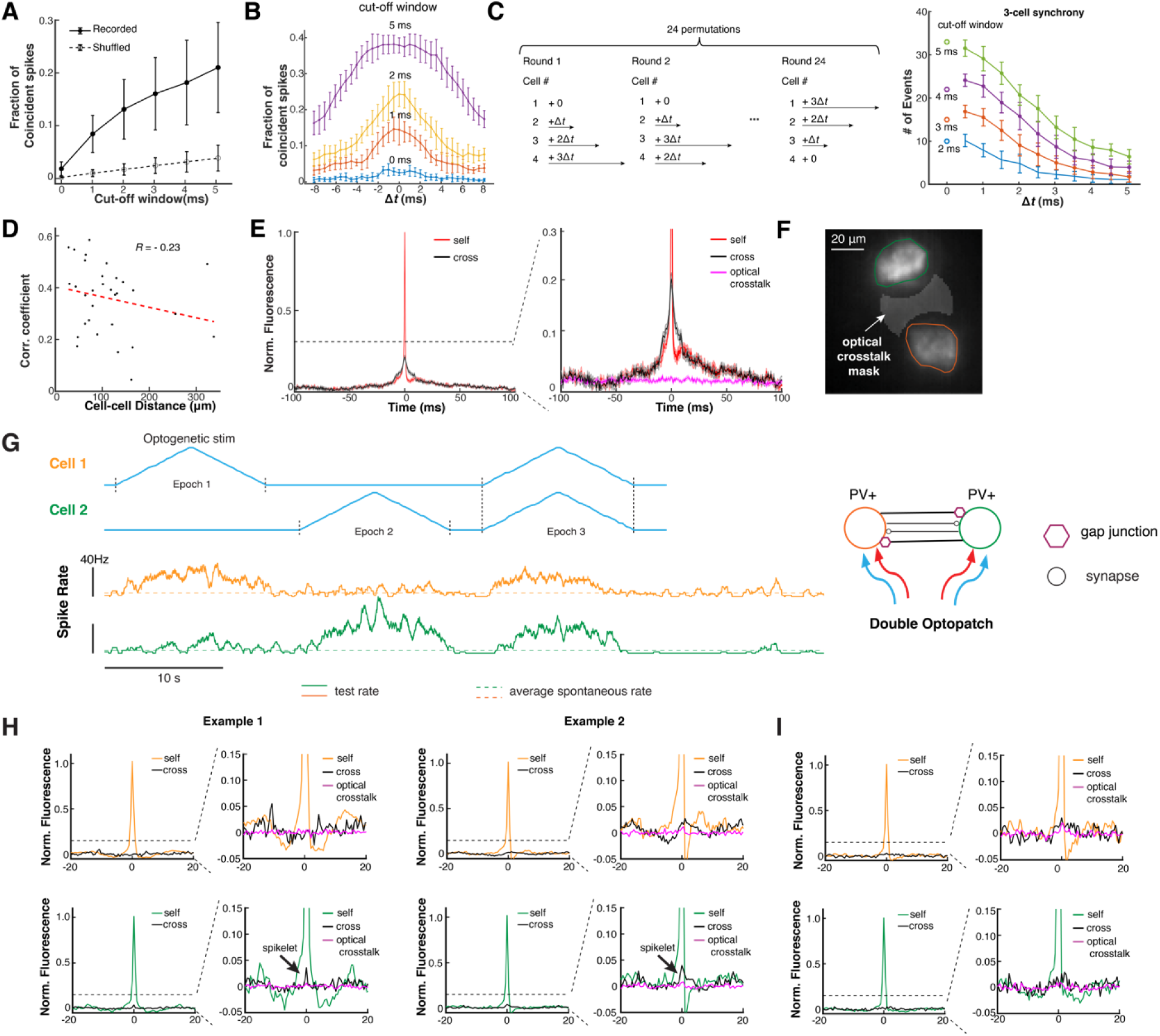
Correlated dynamics between hippocampal PV cells, related to Figure 7. (A) Fraction of synchronous events within different cut-off windows in the experimentally recorded spike train and simulated spike train obtained by random shuffling of the interspike intervals. Error bar: SD. (B) Effect of time-shifting one spike train by Δ*t* on pairwise spike synchrony in six highly correlated pairs of PV cells. The cut-off window indicates the upper limit of the gap between spike maxima. Error bar: SD. (C) Effect of time-shifting the spike trains by multiples of Δ*t* on pairwise and three-way synchrony in the four-cell recording shown in Figure 7C, Error bar: SD. (D) Correlation of subthreshold dynamics vs. inter-soma distance (*n* = 31 pairs from 2 animals). (E) Comparison of self- and cross spike-triggered voltage waveform for events where both cells spiked within 10 ms. (F) Extracting fluorescence traces from cell masks and optical crosstalk mask. (G) Schemes for probing chemical and electric synapses in PV cells. The experiments were done in lightly anesthetized mice where PV cells spiked spontaneously (*n* = 8 pairs from 2 animals). In the example shown here, optogenetic stimulation drove independent spike rate change in each cell. The dashed lines represent the average spontaneous spike rate determined from separate recordings without optogenetic stimulation. (H) Two examples where gap junction-induced spikelets were detected between PV pairs. The inter-soma distances were 42 μm (Example 1) and 90 μm (Example 2). (I) An example where no gap junction-induced spikelet was detected between the PV pair (inter-soma distance = 65 μm).

