## Supplementary material for "All-optical electrophysiology with improved genetically encoded voltage indicators reveals interneuron network dynamics in vivo": Materials and Methods

**Affiliations:**

### MATERIALS AND METHODS

| REAGENT or RESOURCE | SOURCE | IDENTIFIER |
|---|---|---|
| **Bacterial and Virus Strains** | | |
| AAV2/9.hSyn::Cre.WPRE | UPenn Vector Core | N/A |
| AAV2/9.hSyn::DiO:SomQuasAr6a-EGFP-P2A-somCheRiff-HA.WPRE | Janelia Viral Tools | N/A |
| AAV2/9.hSyn::DiO:SomQuasAr6b-EGFP-P2A-somCheRiff-HA.WPRE | Janelia Viral Tools | N/A |
| **Chemicals, Peptides, and Recombinant Proteins** | | |
| doxycycline hyclate | Sigma | Cat# D9891 |
| Blasticidin S | Sigma | Cat# 203350 |
| Puromycin dihydrochloride from *Streptomyces alboniger* | Sigma-Aldrich | Cat# P7255 |
| Geneticin® Selective Antibiotic (G418 Sulfate) | ThermoFisher | Cat# 11811023 |
| HA Tag recombinant rabbit monoclonal antibody | ThermoFisher | Cat# RM305 |
| goat anti-Rabbit IgG (H+L) cross-adsorbed secondary antibody conjugated with Cyanine5 | ThermoFisher | Cat# A10523 |
| **Experimental Models: Cell Lines** | | |
| HEK293T | ATCC | ATCC CRL-3216 |
| tet-on spiking HEK cell | This work | ATCC CRL-3479 |
| CheRiff-EGFP tet-on spiking HEK cell | This work | ATCC CRL-3480 |
| CheRiff-CFP tet-on spiking HEK cell | This work | N/A |
| **Experimental Models: Organisms/Strains** | | |
| C57BL/6 wild-type mice | Charles River | Strain Code 027 |
| NDNF-Cre transgenic mice | Jackson Lab | Stock #028536 |
| PV-Cre transgenic mice | Catherine Dulac | N/A |

| Recombinant DNA | | |
|---|---|---|
| TDG004 pLenti_CMVtight_Kir2.1-CFP | This work | |
| pLenti-CMV-rtTA3-Blast | Eric Campeau | Addgene #26429 |
| HT028 FCMV_CheRiff-EGFP | This work | |
| HT041 FCMV_CheRiff-CFP | This work | Addgene #136636 |
| HT063 FCMV_Archon1-Citrine | This work | |
| HT075 Fsyn_FAS(Cre-off) Archon1-Citrine | This work | |
| HT091 FCMV_Archon1-dark citrine_P2A_mEos4a | This work | |
| HT103 FCMV_QuasAr6a-Citrine | This work | |
| HT110 FCMV_QuasAr6b-Citrine | This work | |
| HT111 Fsyn_FAS(Cre off)_QuasAr6a-Citrine | This work | |
| HT114 Fsyn_FAS(Cre-off)_QuasAr6b-Citrine | This work | |
| HT107 pAAV_hSyn-DiO-SomQuasAr6a_EGFP-P2A-somCheRiff_HA | This work | |
| HT115 pAAV_hSyn-DiO-SomQuasAr6b_EGFP-P2A-somCheRiff_HA | This work | |
| **Software and Algorithms** | | |
| MATLAB R2016b - 2020a | Mathworks | Matlab |
| Labview 2014, 2015 | National Instruments | Labview |
| NoRMCorre | (Pnevmatikakis et al., 2017) | https://github.com /flatironinstitute/NoRMCorre |
| **Others** | | |
| Custom-designed ultra-widefield microscope | (Werley et al., 2017b) | N/A |
| Custom-designed structured illumination microscope | (Fan et al., 2020) | N/A |

### METHOD DETAILS

#### Molecular cloning

Restriction endonucleases were purchased from New England BioLabs (NEB). Non-mutagenic PCR reactions were performed with Phusion® High-Fidelity DNA Polymerase (NEB, Cat. # M0530L). Synthetic DNA oligonucleotides used for cloning were purchased from Integrated DNA Technologies (IDT). Opsin sequences containing a single point mutation were generated through site-directed mutagenesis (QuikChange Lightning Single or Multi kit, Agilent Technologies, Part # 210518 or 210519). Opsin sequences containing multiple point mutations were synthesized *de novo* as gBlocks (IDT). Error-prone PCR was performed with GeneMorph II Random Mutagenesis Kits (Agilent Technologies, Part # 200552) or home-made PCR cocktail (NEB Taq polymerase, 5 mM $MgCl_2$, 0.2 mM each of dGTP and dATP, and 1.0 mM each of dCTP and dTTP). Small-scale isolation of plasmid DNA was performed in house with GeneJET miniprep kit (Thermo Scientific, Cat.# K0503). Large-scale isolation of plasmid DNA was outsourced to Genewiz.

#### HEK cell culture

Wild-type or engineered HEK293T cell lines were maintained at 37 °C, 5% $CO_2$ in Dulbecco's Modified Eagle Medium (DMEM) supplemented with 10% fetal bovine serum, 1% GlutaMax-I, penicillin (100 U/mL), streptomycin (100 mg/mL). For maintaining or expanding the cell culture, we used TC-treated culture dish (Corning). For all the imaging experiments, cells were plated on glass-bottomed dish dishes (Cellvis, Cat.# D35-14-1.5-N). Before optical stimulation and imaging, the medium was replaced with extracellular (XC) buffer containing 125 mM NaCl, 2.5 mM KCl, 3 mM $CaCl_2$, 1 mM $MgCl_2$, 15 mM HEPES, 30 mM glucose (pH 7.3). We found that the XC buffer maintained the cell adhesion and response to optogenetic stimulation for at least 7 - 8 hours.

#### Lentivirus packaging

All the lentivirus preparations were made in house. HEK293T cells were co-transfected with the second-generation packaging plasmid psPAX2 (Addgene #12260), envelope plasmid VSV-G (Addgene #12259) and transfer plasmids at a ratio of 9:4:14. In this study, we generally used FCMV, sometimes pLenti-CMV, as the transfer vector for HEK cell experiments. We used FSyn for cultured neuron experiments (Fan et al., 2018). Both FCMV and FSyn were modified from a previously described FCK lentivirus vector (Hochbaum et al., 2014) by replacing the original CaMKII with a CMV or a hSyn promoter, respectively. For lentivirus intended for HEK cell transduction, 2.7 μg total plasmids for a small culture (300k cells in 35-mm dish) gave sufficient yield of lentivirus. For cultured neuron transduction, larger cultures in 15-cm dish or 10-layer

HYPER Flasks (CheRiff construct, Corning #10030) were used, and HEK cells were transfected with PEI using established protocols (Nguyen et al., 2019). The harvested Virus was concentrated 10-fold (voltage sensors) or 30-fold (CheRiff) using a cationic polymer (Takara Lenti-X Concentrator).

**Photoselection system**

*Optical system*

The optical system was described in an earlier publication (Werley et al., 2017b) with a few modifications for the present use. The microscope was in an inverted configuration to facilitate imaging of cultured cells in glass-bottomed dishes. The system was equipped with several light sources delivered to the sample through free-space optics: 1) a 635-nm laser (DILAS 8 Watts, MB-638.3-8C-T25-SS4.3) sent to the sample plane from below through a near-total internal reflection (near-TIR) configuration for imaging archaerhodopsin-derived GEVIs; 2) LEDs mounted from the above for optogenetic stimulation and imaging fluorescent proteins; 3) a 405-nm laser (MDL-W-405-1W) projected to a micromirror-array device (Digital light innovations, Discovery D4100 with DLP9500 chip and ALP 4.1 High-Speed control software) for photoselection. The patterned light was collected with a tube lens (Olympus MVX, 0.63×) and directed to the sample by a small 45° mirror (4 mm mirror, Tower Optical, MPCH-4.0) inserted into the infinity space. The emission fluorescence was collected with a low-magnification (2×) and high-numerical aperture (NA 0.5) objective lens and filtered with wavelength-specific filters inserted into the infinity space. The emission filter wheel was tilted by a small angle to avoid reflection of light between the sample dishes and filters. After filtering, the emission light was reimaged through a tube lens (Zeiss, Milvus 2/135) and recorded with a scientific CMOS camera (Hamamatsu, ORCA-Flash 4.0). The system was controlled by custom-made LabView codes.

*Calibration of patterned illumination*

The DMD comprised a 1920 ×1080 array, which did not provide 1:1 correspondence with the camera chip (1024×1024 at binning = 2). Moreover, small alignment errors and optical aberrations could manifest as substantial projection errors. To register the DMD array with the camera pixel coordinates, we projected an equally spaced ($d$ = 50 pixels), 11×11 array of dots onto a fluorescent exposure target. The dimension of this test pattern (500 × 500 pixels) was intended to cover the FOV for GEVI screening. The center of each dot in the camera image was determined by 2D Gaussian fitting of the point-spread function. The expected and the observed coordinates of the projection centers were used to construct a piecewise-linear affine spatial transformation.

The resulting projection errors of the 11×11 array were 0.2 - 0.4 pixels (1 - 2 μm), well below the size of mammalian cells. The calibration was performed at the start of each day's experiments and remained stable throughout the day.

*Measuring the optical dose-response curve of phototaggable fluorescent proteins*

mEos4a and PA-mCherry were gifts from Michael Davidson (Addgene plasmid # 54811, #54495). PA-GFP was a gift from Karel Svoboda (Addgene plasmid # 18697). These genes were cloned into FCMV lentiviral vector. Stable HEK cell lines expressing these phototaggable FPs were created by lentivirus transduction.

In the phototransformation experiment, the cells were broadly illuminated from the top with violet light LED (Thorlabs M405L3 + Chroma ET405/20) at an intensity of 0.14 or 0.39 W/cm$^2$. The dark-to-green photoactivation of PA-GFP was monitored in the green channel (Thorlabs M470L3 + Semrock FF01-475/28-25). The green-to-red photoconversion of mEos4a and dark-to-red photoactivation of PA-mCherry were monitored in the red channel (Thorlabs M530L3 + Semrock FF02-529/24-25).

*Selective photoconversion of mEos4a$^+$ cells among PA-mCherry$^+$ cells*

mEos4a$^+$ cells and PA-mCherry$^+$ cells were mixed together at a ratio of 1:20 and plated on the glass-bottomed dish. The cell growth was monitored until a confluent monolayer was formed. A green channel image (Thorlabs M470L3 + Semrock FF01-475/28-25) and a red channel image (Thorlabs M530L3 + Semrock FF02-529/24-25) were taken. A binary mask was generated to isolate the green cells. This binary mask was converted into a DMD mask to address the mEos4a$^+$ cells. Two epochs of 405-nm illumination (25 mW/cm$^2$, 10 min) were successively applied: 1) patterned illumination targeting mEos4a$^+$ cells, followed by 2) wide-field illumination of the entire FOV. After each 405-nm illumination, a red channel image was taken to evaluate the spatial specificity of phototransformation. In the end, the cells were treated with 200 μL trypsin (1%) for 5 min at 37 °C and carefully transferred into 15 mL Falcon tube. The cells were gently centrifuged to remove the trypsin and washed once with the XC buffer. Then the cells were resuspended in the XC buffer and subjected to FACS within one hour.

**Video-based pooled screening for engineering improved GEVIs**

*Engineering monoclonal spiking HEKs*

All the spiking HEK cells were engineered on HEK293T background (ATCC CRL-3216). First, $Na_V$1.5-Puro$^+$ HEK293T cells were generated as previously described (Zhang et al., 2016). The tet-inducible expression system was designed by the Eric Campeau lab and obtained through Addgene. $K_{ir}2.1$-CFP was cloned into the open reading frame of pLenti-CMVtight-EGFP-Neo vector (Addgene Plasmid #26586). pLenti-CMV-rtTA3-Blast (Addgene Plasmid #26429) was used directly to package lentivirus. $Na_V1.5$-Puro+ HEK293T cells were simultaneously infected with pLenti-CMV-rtTA3-Blast and pLenti-CMVtight-$K_{ir}2.1$-CFP-Neo (TDG004). The cells were first selected with three antibiotics (2 μg/mL puromycin, 5 μg/mL blasticidin, 200 μg/mL Geneticin/G418), then induced with doxycycline (2 μg/mL) for ~30 hours before FACS purification. The $CFP^+$ cells were seeded into 96-well plates (1 cell per well) and cultured 3 - 4 weeks under the standard conditions for HEK cell culture. The expanded monoclonal cells were screened with current clamp. The clone that showed robust spike upon current injection was termed tet-on spiking HEK cell and used to engineer the CheRiff-$CFP^+$ spiking HEK cells.

CheRiff-CFP was cloned into FCMV lentivirus vector (HT041). After lentiviral infection and monoclonal selection, the CheRiff-$CFP^+$ spiking HEK cells were optically screened. The spikes were evoked with optogenetic stimulation (ex = 490 nm) and visualized using a voltage-sensitive dye (BeRST, ex 635 nm) (Huang et al., 2015). The CheRiff-$EGFP^+$ spiking HEK cells were engineered differently. Nav1.5-Puro/rtTA3-Blast/$K_{ir}2.1$-CFP-$Neo^+$ polyclonal HEK cells were infected with FCMV-CheRiff-EGFP (HT028) lentivirus. After doxycycline induction, the $CFP^+$/$EGFP^+$ cells were purified by FACS and seeded into 96-well plate for monoclonal selection. The monoclonal cells were validated by patch clamp under optogenetic stimulation.

To enhance genomic stability, the spiking HEK cells can be maintained in antibiotic-containing medium (2 μg/mL puromycin, 5 μg/mL blasticidin, 200 μg/mL Geneticin/G418). However, we found the cell lines reasonably stable even without these antibiotics. To obtain consistent experimental results, we only used low passage-number cells and kept a master plate free of doxycycline.

Tet-on spiking HEK cell and CheRiff-$EGFP^+$ tet-on spiking HEK cell are available from ATCC (CRL-3479; CRL-3480).

*Generation of the library cells*

Archon1 sequence was a gift from Ed Boyden at MIT. Previously, we found that a fluorescent protein tag could significantly enhance the membrane localization of Archaerhodopsin-derived GEVIs in mammalian cells. In particular, a combination of Citrine and multiple repeats of trafficking sequence (TS-Citrine-TS×3-ER2) improved the voltage imaging SNRs in cultured neuron (Adam et al., 2019). Initially, we attempted to substitute Citrine with mEos4a. However, this substitution resulted in poorly trafficked protein. Therefore, we switched to a bicistronic construct, in which GEVI and mEos4a were linked with a self-cleaving P2A peptide (Kim et al., 2011). Because Citrine and mEos4a share the same spectral window, a single point mutation (Y67G) was introduced into Citrine to create a non-fluorescent protein tag, "dark Citrine" (Fan et al., 2018).

Random mutations were introduced into Archon1 using error-prone PCR. Then the mutated opsin sequences were fused to the rest of the coding sequence (TS-dark Citrine-TS×3-ER2-P2A-mEos4a) using fusion PCR and purified with agarose gel electrophoresis. The purified DNA fragment was inserted into the FCMV lentivirus vector using Gibson assembly, transformed into DH5$\alpha$ *E.* coli competent cells (NEB), and plated on ampicillin-containing (Amp+) agar plates. We used Sanger sequencing to analyze the mutation rate of a small number (<10) of clones. Each mutant included 0, 1, or 2 amino acid substitutions (average number ~ 1). The colonies were scraped from the agar plates, transferred into Amp+ LB medium, allowed to grow at room temperature for approximately 1 hour before miniprep. The plasmid library was then used for lentivirus preparation. CheRiff-CFP$^+$ spiking HEK cells were infected with the lentivirus library at a low titer (MOI ~ 0.01). The mEos4a$^+$ cells were purified with FACS and cultured with the standard HEK cell culture protocol.

*Screening*

The library cells were mixed with CheRiff-CFP$^+$ spiking HEK cells (spacer cells) at a ratio of 1:10. The mixed cells were plated in glass-bottomed dishes (Cellvis, Cat. # D35-14-1.5-N) homogenously at a density of 500k cells in a 14-mm well. The cells were allowed to grow for 40-50 hours in Dox$^+$ medium to form a monolayer. Before imaging, the medium was replaced with XC buffer.

Before each screening experiment, the DMD projection was recalibrated as described above. To achieve near-TIR excitation, the gap between the well and the prism window was filled with index-matching immersion oil (Olympus, Z-81114). A 500×500 pixel (binning = 2) field of view (FOV) was used for GEVI characterization. First, an mEos4a-channel image was taken (excitation: 490-nm LED, Thorlabs, M490L3; emission filter: 540/50 Semrock FF01-540/50). Next, the spiking HEK cell monolayer was broadly stimulated with the 490-nm pulses and the voltage responses to optogenetic stimulations was recorded (excitation: 635-nm laser, 100 W/cm$^2$; emission filter: long-pass 700 nm; sampling rate:100 Hz). Precisely timed spikes were evoked by blue light (490 nm, 10 ms) stimulation from the top. In each experiment run, the cells were stimulated with 5 blue light pulses with the red laser was on, then stimulated with 5 blue light pulses while the red laser off. The red laser-off epoch was used to correct for blue light crosstalk.

The 500 × 500-pixel FOV (2.3 mm × 2.3 mm in the sample plane) was segmented based on the mEOS-channel (excited with 490-nm LED) image using custom Matlab code. Briefly, the mEos4a-channel image was corrected for baseline and segmented with Watershed algorithm. Only ROIs larger than 5 pixels were accepted as "cells", and smaller ROIs were rejected. We observed that after 2 days of culture, the library cells often showed small clusters, primarily due to cell division. We reasoned that since the neighboring cells descended from the same parent cells, it was acceptable to treat small clusters as a single genotype.

The Arch-channel movie was corrected for blue-light crosstalk, red-light excitation profile, and baseline. The ROIs generated based on the mEos4a-channel was mapped onto the movie. The average intensity traces were extracted from each ROI and corrected for photobleaching. The initial fluorescence intensity was assigned as $F_0$. The averaged baseline-to-peak difference was assigned as $\Delta F$. SNR was estimated as $\Delta F_{\mathrm{Arch}} / \mathrm{sqrt}(F_{0,\mathrm{Arch}})$.

In pilot experiments on monoclonal CheRiff$^+$/Archon1-Citrine$^+$ spiking HEK cells, we found that even a monoclonal population showed substantial variation in $F_0$ and $\Delta F$, possibly due to variations in protein expression or trafficking. Thus, we concluded that 1) there were non-genetic factors underlying the broad distribution of expression level, and 2) setting a stringent threshold for $F_0$ was unlikely to be meaningful. In our experiments, we set a 50$^{\mathrm{th}}$ percentile threshold for $F_0$, and 75$^{\mathrm{th}}$ percentile cut-off for $/ \mathrm{sqrt}(F_{0,\mathrm{Arch}})$. These top 12.5% of ROIs were selectively illuminated with violet light to create a binary marker.

A DMD mask was generated to address the selected ROIs. The photoconversion for each FOV took 10 min. Then the dish was moved to allow the next FOV to be characterized in the same manner. The spiking HEK cells responded robustly to optogenetic stimulation throughout a time course over 6 hours. In a typical experimental run, 20 - 25 FOVs were scanned to achieve good coverage of the entire dish, and each FOV contained 2,000 - 4,000 ROIs. From a single dish, 30,000 to 50,000 cells were scanned.

The cells were then treated with 200 μL trypsin (1%) for 5 min at 37 °C and carefully transferred into 15 mL Falcon tube. The cells were gently centrifuged to remove the trypsin and washed once with the XC buffer. Then the cells were resuspended in the XC buffer and subjected to FACS within one hour. The photoconverted cells were collected into fresh DMEM10 medium and cultured under the standard HEK cell culture condition to expand the population. In each round of enrichment, 2 dishes (approximately 90k library cells) were screened. 7 - 10 days later, the enriched library from the 2 dishes were combined at a proportion corresponding to the number of originally collected cells and subjected to the next round of enrichment. The remaining library cells were preserved in liquid nitrogen for sequencing.

*Illumina sequencing*

16k - 20k library cells were collected into PCR tubes and boiled (98 °C, 5 min) to release their genomes as the PCR template. The opsin sequences were amplified from the genome (forward primer: GACCTCCTCGGAGATGGTAGA; reverse primer: AGCTGAAGGTTCAGGTGCTTC). The primer pair used here gave a 720-bp amplicon that covered the 31-750 nt of Archon1 CDS, which effectively provided single nucleotide polymorphism (SNP) information for 52-729 nt. We also attempted primers designed to cover the entire 759 nt of Archon1 CDS. However, we found that full-coverage primers did not result in robust PCR amplification. We reasoned that as the N-terminus and C-terminus are distant from the retinal chromophore and unlikely to modulate the voltage-sensitive fluorescence, the omission of the terminal sequence information should not severely negatively impact our screening efforts. In the earlier efforts to optimize archaerhodopsin-derived GEVIs, no beneficial mutations have been identified in the missing regions (Hochbaum et al., 2014; Piatkevich et al., 2018).

The 720-bp amplicon were then segmented into 3 smaller amplicons with high-fidelity PCR (Fwd1: TCGTCGGCAGCGTCAGATGTGTATAAGAGACAG-GACCTCCTCGGAGATGGTAG, Rev1: GTCTCGTGGGCTCGGAGATGTGTATAAGAGACAG-TGTAGTGAACAGCCACTGTG; Fwd2: TCGTCGGCAGCGTCAGATGTGTATAAGAGACAG-CTGAACATCTACTACGCAAG, Rev2: GTCTCGTGGGCTCGGAGATGTGTATAAGAGACAG-CTGGGCCTCTCTCCTTAGCG; Fwd3: TCGTCGGCAGCGTCAGATGTGTATAAGAGACAG-GTCCTGGCCACTTCTCTGCG; Rev3: GTCTCGTGGGCTCGGAGATGTGTATAAGAGACAG-AGCTGAAGGTTCAGGTGC). We chose 2 x 150 bp paired-end MiSeq (Harvard Medical School Biopolymer Facility) to analyze the SNPs. We obtained a sequencing depth of 2 - 5 $\times 10^5$ reads per nt (filtered for Illumina Q score > 30). VCF data were generated from the FASTAQ data with a custom pipeline that included Trimomatic (Bolger et al., 2014), NGmerge (Gaspar, 2018), BWA (Li et al., 2009a), samtools (Li et al., 2009b), and Pilon (Walker et al., 2014). The VCF data were subsequently analyzed with custom Matlab code.

*Simulation of the selection threshold*

To determine the probability that a mutation could be enriched in the selection by chance alone, we performed a Matlab simulation of the selection process, assuming random selection for a mutant with starting frequencies of 0.002%, 0.004%, 0.006%, 0.008%, 0.01%, 0.02%, 0.03%, 0.04%, 0.05%, 0.06%,0.07, 0.08%. The upper limit of 0.08% was chosen based on the sequencing result of the starting library. The initial number of library cells was set to be 50,000. We randomly allowed 5,000 – 12,000 “cells” to pass the selection. These numbers were determined by the actual numbers of cells collected from FACS (5,000 ~12000 cells after each round). We then expanded the population 100-fold and repeated the sampling and expansion process two more times. The 95% confidence threshold on the prevalence of a mutational frequency arising by chance was determined from 2,000 iterations of the simulation.

**Characterization of improved GEVIs in cell culture**

*Imaging and electrophysiology of HEK293T cells*

To prepare the samples for characterization in HEK cells, the GEVI constructs with the appropriate mutations were cloned into FCMV vector (HT63, HT103, HT110) and packaged into lentivirus. HEK cells we infected at a low-titer (MOI < 0.1) and purified by FACS.

All imaging and electrophysiology experiments were performed in XC buffer. Concurrent whole-cell patch-clamp and high-magnification fluorescence recordings were acquired on a custom-built, dual-view, inverted epifluorescence microscope equipped with the electrophysiology module described before (Adam et al., 2019). For fluorescence measurement, a high-magnification water-immersion objective (Olympus, 60×, NA 1.2) was used. The GEVI fluorescence was excited by a 635 nm laser (420 $W/cm^2$), filtered with a dichroic (Semrock; FF640-FDi01-25×36) and a Cy5-longpass filter (708/75, and imaged with a sCMOS camera (Hamamatsu, ORCA-Flash 4.0). The citrine fluorescence was excited with 488 nm laser (100 - 200 $mW/cm^2$), filtered with a GFP filter (Semrock 525/30), and imaged with an EMCCD camera (Andor iXonEM+ DU-897E). Filamented glass micropipettes were pulled to a tip resistance of 5 - 8 MΩ, and filled with internal solution containing (in mM): 125 potassium gluconate, 8 NaCl, 0.6 $MgCl_2$, 0.1 $CaCl_2$, 1 EGTA, 10 HEPES, 4 Mg-ATP and 0.4 Na-GTP (pH 7.3); adjusted to 295 mOsm with sucrose. Pipettes were positioned with a Sutter MP285 manipulator. Whole-cell patch clamp recordings were performed with a MultiClamp 700B amplifier (Molecular Devices), filtered at 2 kHz with the internal Bessel filter and digitized with a National Instruments PCIE-6323 acquisition board at 10 kHz. For photocurrent measurement, the 635 nm laser intensity was 1500 $W/cm^2$ and the 488-nm laser intensity was 124 $W/cm^2$.

*Imaging and electrophysiology of cultured rat hippocampal neuron*

To prepare the samples for characterization in HEK cells, the GEVI constructs with the appropriate mutations were cloned into a neuronal expression lentiviral vector under hSyn promoter (HT111, HT114). The neurons were transfected via the calcium phosphate method or transduced with lentivirus on DIV7 and used for imaging or electrophysiology experiments between DIV11 - DIV14.

The optical system for concurrent fluorescence and electrophysiological recordings in cultured neurons was the same as in the HEK293 cell experiments. The internal solution for whole-cell patch clamp contained (in mM), K-gluconate 135, KCl 4, HEPES 10, EGTA 0.5, $Na_2$-phosphocreatine 10, Mg-ATP 4, and $Na_2$-GTP 0.4, 292-300 mOsm, pH 7.3.

*High-throughput imaging of hippocampal neurons*

Primary E18 rat hippocampal neurons (fresh, never frozen, BrainBits #SDEHP) were dissociated following vendor protocols and plated in PDL-coated 96-well plates. Neurons (21k/$cm^2$) were cocultured with primary rat glia (27k/$cm^2$) to improve cell health and maturation. Custom 96-well plates from ibidi GmbH had the standard low-absorption, low-autofluorescence cyclic olefin copolymer (COC) foil substrate and clear COC walls to minimize laser absorption. Lentivirus for Archon1-EGFP, Archon1-Citrine (HT075), QuasAr6b-Citrine (HT111), QuasAr6b-Citrine (HT114) were packaged in parallel under identical conditions. The virus titers were confirmed with qPCR. Neurons were transduced after 6 days in culture with 1) 0.33 µL lentivirus encoding CheRiff-EBFP2 driven by the synapsin promoter and 2) varying doses of the voltage sensor variants, also driven by the synapsin promoter. Functional Optopatch imaging was performed after 14 days in culture.

Imaging was performed on the Firefly microscope. Optogenetic stimulus to CheRiff was generated by a blue LED, filtered (Semrock No. FF01-470/28), and delivered to a large area with intensity ranging from 2 to 88 mW/$cm^2$. 638 nm red laser light was applied through a prism in near-TIR, so the beam transmitted into the imaging media and propagated nearly parallel to the surface. The illumination intensity was 200 W/$cm^2$ (neglecting beam intensification by refraction at the imaging buffer/COC substrate). Fluorescence was imaged at 2.7× magnification onto an sCMOS camera (Hamamatsu ORCA-Flash 4.0 V2) through a near-infrared emission filter (Semrock #FF02-736/128) and data was collected at a 1 kHz frame rate.

The movie was segmented and detected using proprietary Matlab code developed by Q-State Biosciences. Briefly, the spiking neurons were automatically detected and segmented using a principal component analysis/independent component analysis (PCA/ICA)-based Matlab code described in (Werley et al., 2017a). The algorithm identifies spatially compact sets of pixels (neuron masks) that co-vary in time with action potential positive-going voltage transients. A quality control process discards sources whose action potential height does not exceed the baseline noise by at least a factor of 3. Improved voltage sensor SNR, membrane trafficking, or expression level raises the signal of more sources above the noise floor, leading to automatic identification of more spiking neurons per field of view.

*Confocal imaging of QuasAr6a and QuaAr6b expressed in cultured rat hippocampal neuron*

To sparsely expression QuasAr6b-Citrine or QuasAr6b-Citrine, rat hippocampal neuron cultures were transfected with Fsyn plasmids (HT111, HT114) encoding the constructs via Ca-Phos. Before imaging, the medium was replaced with transparent XC buffer. The confocal images were acquired on LSM880 Airyscan with an air 20× objective. Citrine fluorescence was excited with 488-nm laser.

**Characterization of QuasAr6a and QuasAr6b in brain slice**

*Expression of the Optopatch constructs in tissue*

The design of the cre-on bicistronic construct of QuasAr6b and QuasAr6b (pAAV_hSyn-DiO-SomQuasAr6-EGFP-P2A-somCheRiff_HA) was based on Optopatch4 that used Archon1 as the voltage indicator (Fan et al., 2020). The high-titer Optopatch viruses (AAV2/9 hSyn-DiO-SomQuasAr6a-EGFP-P2A-somCheRiff-HA; AAV2/9 hSyn-DiO-SomQuasAr6b-EGFP-P2A-somCheRiff_HA; AAV2/9 hSyn-DiO-SomArchon1-EGFP-P2A-somCheRiff-HA) were obtained from the Janelia Vector Core. Before the surgery, the virus was freshly diluted in PBS to the specified final titer. Expression in cortex was achieved through intracranial injection of AAVs ($5\times10^{12}$ GC/mL Optopatch + $10^{11}$ GC/mL hSyn_Cre) in P0 - P2 wild-type C57BL/6J pups. Specifically, cryo-anaesthetized pups were injected in the left hemisphere, 1.0 mm lateral and 1.0 mm anterior to lambda, starting from a -1.0 mm depth. Diluted virus (40 nL, 60 nL/min) was injected at 0.1 mm increment as the pipette was withdrawn. The pups were recovered on warm heating blanket and returned to the dam.

*Slice preparation and confocal imaging*

Coronal slices were prepared with the injected pups 15 - 21 days after virus injection. The slices were fixed in 1% paraformaldehyde for 3 - 4 hours and immunostained to visualize the HA tag (primary antibody: HA Tag recombinant rabbit monoclonal antibody, ThermoFisher, RM305, 2,000× dilution; second antibody: goat anti-Rabbit IgG (H+L) cross-adsorbed secondary antibody conjugated with Cyanine5, ThermoFisher, A10523, 500× dilution).

The mounted slices (VECTASHIELD® Antifade Mounting Medium H-1000, Vectorlabs, H-1000-10) were imaged on LSM880 Airyscan with 488 nm excitation for the EGFP fluorescence and 635 nm excitation for the Cy5 fluorescence.

**In vivo all-optical electrophysiology**

*Cranial window surgery for imaging visual cortex L1*

For experiments in Figure 4, the window was comprised of two 3-mm round #1 cover glasses and one 5-mm round #1 cover glass (Harvard apparatus) cured together with UV curable adhesive (Norland Products, NOA 81). For experiments in Figure 5, one 3-mm round #1 cover glass was glued to a custom-made stainless-steel adapter. The adapter has an outer diameter of 5 mm and inner diameter of 2.7 mm.

The cranial window surgery for imaging L1 cortex was performed as described previously (Goldey et al., 2014). In brief, 10 - 16 weeks-old NDNF-Cre$^{+/-}$ mice (male or female) were induced with > 2% isoflurane and maintained in deep anesthesia with 1% isoflurane throughout the surgery. A heating pad (WPI, ATC2000) was placed beneath the mice to main the body temperature at 36 - 37 °C. Ophthalmic eye ointment was applied on the eyes to keep them moist. An approx. 3-mm craniotomy was created on the left visual cortex of the exposed skull (AP: 2.5 - 2.6-mm lateral, 0.8-mm anterior of lambda) with a dental drill. The Optopatch virus was diluted to a final titer of $1\times10^{13}$ GC/mL for experiments in Figure 4, or to a final titer of 0.5 - $1.0\times10^{13}$ GC/mL for experiments in Figure 5. The diluted virus was injected at 7 - 8 sites across the craniotomy (80 and 160 µm beneath dura; 40 - 60 nL each depth; 30 - 60 nL/min). After virus injection, the craniotomy was covered with the glass window. The edge of the window was glued to the skull with cyanoacrylate adhesive (3M Vetbond). Next, a titanium headplate (designed based on (Goldey et al., 2014)) was attached to the exposed skull with dental cement (C&B metabond, Parkell, No. 242-3200). Special care was taken to ensure that the dental cement filled the space between the rim of the window and the skull and covered all the exposed area of the skull. Animal typically recover from anesthesia within 20 min. Then they were returned to their home cage and administrated with Carprofen (5 mg/kg) and Buprenorphine (0.1 mg/kg) on post-surgery Day 0, 1, 2.

*Window surgery for imaging hippocampus CA1*

The window surgery for imaging hippocampus CA1 was performed based on previous reports (Adam et al., 2019; Dombeck et al., 2010). In brief, the cannula window was comprised of a 1.5-mm segment of a 3-mm outer diameter thin-walled stainless steel tube (MicroGroup) and 3 mm

#1 round cover glass (Harvard Apparatus) glued to one end of the tube using UV-curable adhesive (Norland Products, NOA 81). 8 - 12 weeks old PV-Cre$^{+/-}$ mice (male or female) were used for imaging. A 3-mm craniotomy was created on the left hemisphere (1.8 mm lateral, 2.0 mm caudal of bregma) with a biopsy punch (Miltex). Optopatch virus was diluted to 2.5 ~ 5 $\times 10^{12}$ GC/mL and injected into three sites near the center of the craniotomy (1.0 mm to 1.4 mm beneath dura with 0.1 mm increment; 40 nL each depth; 60 nL/min). After virus injection, the cortex was carefully aspirated, and the center region of the external capsule was removed to expose the hippocampus CA1. The cannula was then lowered onto the CA1 surface until the window touched the tissue. The remaining outer surface of the cannula was sealed to the exposed skull with dental cement (C&B Metabond). Finally, a titanium head plate was fixed onto the exposed skull. The post-surgery care was identical to that of the cranial window surgery for L1 imaging.

*Animal handling for in vivo imaging*

Head-fixed animals were imaged in various degrees of anesthesia or full wakefulness. For imaging experiments under deep anesthesia, 1 - 1.5% isoflurane was supplied, and the dose was adjusted throughout the course of the imaging session to maintain a stable breathing rate. For imaging experiments under light anesthesia, animals were first administrated with chlorprothixene (0.2 mg/ml, 5 μL/g weight mouse). In the imaging session, 0.4 - 0.7% isoflurane was supplied to keep the animal in a state of semi-wakefulness, with occasional body movements. In all experiments involving anesthesia, the animal was kept on a heating pad (WPI, ATC2000) to maintain stable body temperature at 37 °C, and their eyes were kept moist using ophthalmic eye ointment. A typical imaging session lasted 1 - 2 h, after which the animal generally recovered within 5 min. For imaging experiments under full wakefulness, the animal was first habituated to head restraint in a body tube prior to the imaging sessions, and no extra heating was necessary.

*Optical systems for in vivo all-optical electrophysiology*

The imaging set-up was originally described in (Fan et al., 2020) with a few modifications. For the red laser path, the 639 nm laser source ((CNI Lasers, MRL-FN-639, $\lambda$ = 639 nm, 700 mW single transverse mode) was attenuated with a half-wave plate and polarizing beam splitter, expanded to a collimated beam of ~10 mm diameter, then projected onto the surface of a reflection-mode liquid crystal spatial light modulator (SLM, Meadowlark 1920SLM VIS) with a resolution of 1920×1152 pixels. For the blue laser path, the blue laser (Cobolt, 06-01 series, $\lambda$ = 488 nm, 60 mW) was modulated in intensity via an acousto-optic tunable filter (AOTF; Gooch and Housego TF525-250-6-3-GH18A) and collimated to a beam of ~17 mm in diameter before being directed onto the reflective surface of a digital micromirror device with a resolution of 1024×768 pixels

(DMD, Vialux, V-7001 VIS). The waveforms of optogenetic stimulation sequence and voltage imaging sequencing were controlled via a National Instruments DAQ (NI PCIe-6363). The movies were acquired at 1,000 - 4,000 Hz with a sCMOS camera (Hamamatsu ORCA-Flash 4.0). A Cy5 emission filter was used in the Arch-channel. The camera's internal 100 kHz clock was used as the master clock to synchronize all the DAQ inputs and outputs. The system was controlled with a custom software developed in Matlab. This Matlab-based control software includes modules interfaced with 1) the sCMOS camera, 2) DAQ; 3) DMD, and 4) SLM. The software also includes routines for registration of SLM, DMD and camera.

*In vivo imaging conditions*

For experiments in Figures 4, 5, 6, imaging was performed with a 25× water immersion objective (Olympus XLPLN25XWMP2) with a 2-mm working distance and a numerical aperture of 1.05. For experiments in Figure 7, a 10× water immersion objective (Olympus XLPLN10XSVMP) with an 8 mm working distance and a numerical aperture of 0.6 was used to obtain a larger FOV. To ensure stable water interface between the window and the 10× objective, a 3D-printed adapter hat was attached to the headplate temporarily with vacuum grease during the imaging session.

For voltage imaging, red laser excitation was targeted to the cell membrane or whole soma with holographic optics. In the experiments in Figure 4 where the SNR and kinetics between QuasAr6a, QuasAr6b and Archon1 were compared in $NDNF^+$ cells, 5 mW of red light was targeted to the membrane of the soma. In the experiments in Figure 5 where the monosynaptic connections between $NDNF^+$ cells were measured, 3 - 4 mW of red light was targeted to each cell. In the experiments in Figure 6 the SNR and kinetics between QuasAr6b and Archon1 were compared in $PV^+$ cells, membrane-localized illumination with 10 mW red light was used for each cell. In experiments in Figure 7 where multiple $PV^+$ cells were imaged, 7 - 8 mW red light was used in each cell. In our experiences, membrane-localized illumination gives better SNR when the cells are still. When the cells were experiencing stronger movement, whole-soma illumination helped to reduce motion artifacts.

*Optogenetic stimulation*

For optogenetic stimulation, blue light was patterned to the soma with DMD. The structural image in the GFP channel was excited with a low level of blue light (< 1 mW/mm$^2$) and imaged with a GFP emission filter. The pixel bitmap containing the ROI masks was created based on the GFP channel image. When the experiments required the blue light intensity to change globally for all the ROIs, the blue intensity was modulated with an AOTF upstream of the DMD, with a range from 0 to 25 mW/mm$^2$. When the experiments involved different blue light waveforms for different ROIs (e.g., experiments in Figure 5), the intensity was controlled by randomly switching on a fraction of pixels within the ROI. The pre-defined sequence of pixel bitmaps was loaded into the on-board RAM on the DMD and timed with digital pulses sent from DAQ to the DMD.

For the double Optopatch experiments on NDNF$^+$ and PV$^+$ cells, the stimulation intensity was modulated by randomly switching on a fraction of DMD pixels within each cell mask. For each ramp stimulation, a series of DMD masks were generated and displayed on the DMD as a movie. By varying the fractions of "on" pixels independently for each cell mask, we could achieve different optogenetic stimulation waveforms for different cells.

**Data analysis**

Data were analyzed and plotted with homemade code written in MATLAB.

*Extracting the voltage-sensitive fluorescence*

Movies were corrected for motion using the NoRMCorre algorithm (Pnevmatikakis and Giovannucci, 2017). Next, photobleaching was corrected using mono-exponential fit. Next, We manually created masks for each cell and divided movie into sub-movies based on the contour of the cell masks. To accurately extract the subthreshold dynamics, we performed activity-based image segmentation separately in each sub-movie. Our assumption was that while subthreshold voltages could be correlated between a cell and out-of-focus background cells, spike dynamics are unlikely to be correlated with background. We also assumed that spiking dynamics and the true subthreshold dynamics would share the same spatial footprint. The sub-movies were filtered in time with a 50 Hz high-pass filter, and then segmented semi-automatically using principal components analysis followed by time-domain independent components analysis (PCA/ICA). The spatial masks from PCA/ICA were then applied to the original movies without high-pass filtering to extract fluorescence traces that included both spike and subthreshold dynamics. We found that the quality of the spatial masks generated by PCA/ICA depended on the SNR of the raw movie, as well as the number of spikes in the raw movie. In general, the stronger SNR in the raw movie, and the more spikes in the segmented epoch, the more likely we obtain high-quality spatial masks. For some recordings where the SNR was good enough for accurate detection of spikes (SNR > 4), but did not gave high-quality PCA/ICA masks, these recordings were only used to analyze spike dynamics, but not for extracting subthreshold dynamics.

*Spike detection and trace normalization*

Fluorescence traces were first high-pass filtered at 50 Hz. We used two complementary methods for spike detection. First, a simple threshold-and-maximum procedure was applied on the high-pass filtered fluorescence trace. The initial threshold was set at 3 times of the noise level and adjusted if necessary. Second, we performed wavelet transformation on the high-passed filtered traces to separate the signals based on the time-domain. We next projected the higher-frequency wavelets into a time trace, then applied a threshold-and-maximum procedure to identify the peaks in the projected traces. Compared to Fourier transformation, wavelet decomposition allows the expansion of signals in terms of finite time functions, which gives higher selectivity to impulse-like events such as action potentials. A fluorescence impulse was accepted as spikes only if it stood out in both spike detection methods. We found the wavelet transformation particularly helpful to spike detection in PV neurons because PV neurons' characteristically narrow spikes were much faster than most sources of noises. All fluorescence traces were then normalized to spike height for spike triggered average.

*Analysis of the subthreshold dynamics in PV recordings*

Following spike detection and trace normalization, the frame containing the spike maxima and surrounding the spike maxima (-2.5 ms to + 2.5 ms) were digitally removed. The gap was replaced with the mean of fluorescence signal at -3 ms and + 3 ms. The fluorescence trace was then filtered by applying a moving average filter (moving window = 10 ms). We next adjust the absolute value of the entire traces to set the average of subthreshold signals to zero. We found that even after NoRMCorre motion correction and PCA/ICA segmentation, there could be residual motion artifact, presumably due to z-motion. Thus, we used the NoRMCorre algorithm to generate lateral shift metrics to estimate animal movement. The episodes showing pronounced movement were excluded from the subthreshold analysis. The correlation coefficient of subthreshold dynamics was calculated with Matlab function xcorr. However, we found that even when motion artifact was strong, the spike detection still worked robustly as PV spikes were much faster than any motion artifact. Thus, no episodes were excluded from the spike train.

*Calculation of spike SNRs and waveforms*

We define SNR as the ratio of the height of spike (fluorescence signal above the subthreshold) to the high-frequency noise. The high-frequency noise was defined as the standard deviation of the non-spiking epoch of the high-pass filtered (50 Hz) fluorescence trace. Because the intensity of photocurrent modulated the height and waveform of the spike, we only used the ramp epoch to calculate spike height and waveforms. For PV spikes, we used all the spikes from the upward- and downward- ramps. For the NDNF spikes, because the ramp of the blue light was steeper, we found the spike height and waveform varied quickly. Thus, we only used the first three spikes from the ramp for the calculation.

*Estimation of spike rate with Bayesian Adaptive Kernel Smoother*

In the double Optopatch experiments on the NDNF+ cells, the duration of the stimulation was short so the total number of the spikes were limited. As a result, the calculation of the spike rate was sensitive to the choice of integration window. Thus, we used a with Bayesian Adaptive Kernel Smoother (BAKS) (Ahmadi et al., 2018) to convert discrete spike raster into continuously varying spike rate. The Matlab function was downloaded from https://github.com/nurahmadi/BAKS. The shape parameter (a) and the scale parameter (b) were both set to 40. As a quality control, we compared the average of the BAKS-derived spike rate against the average spike rate directly calculated from the spike raster. We found the two methods in good agreement.

*Calculation of PV spike synchrony*

The cut-off window was defined as the maximal time lag between spiking in two different cells. The number of synchronous spikes was defined as number of incidents where two spikes fell into the cut-off window. The fraction of coincident, or nearly synchronous spikes, was defined as the number of synchronous spikes as a fraction of the average total number of spikes of the cell pair.

*Estimate of optical crosstalk between PV pairs*

We reasoned that the upper limit of optical crosstalk between cell pairs could be estimated using the fluorescence signal extracted from an intervening mask midway between the two cells. The intervening mask was created as follows. First, the area and centroid of the cell masks was determined with Matlab function regionprops. Second, a circular mask was created, with its center coordinates at the mid-point between the centroids of the two cell masks, and its area set to be the sum of the areas of both cell masks. However, when the cells were too close, the intervening mask may overlap with the cell mask. Thus, the pixels located within the circular mask radius from the cell mask centroid were removed from the middle mask. The resulting masks generally had comparable, or larger size than the cell masks. The fluorescence signal was extracted by applying the intervening mask to the motion-corrected movie, corrected for any photobleaching and high-pass filtered, and normalized with the voltage fluorescence traces. The optical crosstalk was calculated as the average fluorescence waveform triggered by the same set of spikes used in calculating cross-triggered average.

*Simulation of PV spike train: shuffling of interspike intervals*

In Figure S7A, the shuffled spike train was created by randomly permutating the interspike intervals. The timing of the first spike in the recording was not perturbed. Then the fraction of coincident spikes for different cut-off windows was calculated for the shuffled spike trains from both cells. For each cell pair, this process was iterated for 20 times. The mean and SEM were calculated from the 20 iterations.

**Statistics**

Statistical tests were performed using standard MATLAB functions (MathWorks). For two-sample comparisons of a single variable, a two-tailed Student's t-test was used when the sample size was large (high-throughput Optopatch in cultured neurons). For datasets where the sample size was small ($n < 40$) or showed non-Gaussian distribution, Wilcoxon rank-sum test was used instead. When calculating the in *vivo* GEVI metrics (Figure 4E, S4B, 6C, D), outliers (value that is more than three scaled median absolute deviations) were excluded. The experiments were not randomized, and the investigators were not blinded to the experimental condition. Sample size was based on reports in related literature and was not predetermined by calculation.  Recordings of non-spiking neurons were excluded from analysis.
